# Cell fusion reprograms tumor cells and promotes RUNX1-mediated invasion and dissemination in colorectal cancer

**DOI:** 10.64898/2026.04.12.717781

**Authors:** Ashley N. Anderson, Konstantin Queitsch, Nicole R. Giske, Jocelyn A. Jones, Amara Pang, Amanda Zucker, Ge Huang, Cody C. Rounds, Brady Jin Smith, John R. Swain, Abigail G. Moore, Divya Ravi, Isis Diaz Monarrez, Sandhya Govindarajan, William S. Greer, Peter H. Tran, Kai Tao, Luiz Bertassoni, V. Liana Tsikitis, Brian Brinkerhoff, Charles D. Lopez, Cristiane Miranda Franca, Jared M. Fischer, Guanming Wu, Young Hwan Chang, Andrew C. Adey, Summer L. Gibbs, Melissa H. Wong

## Abstract

Metastasis remains the primary cause of cancer-related morbidity and mortality, despite significant advances in targeted therapies. Although metastatic dissemination requires tumor cells to escape the primary lesion and colonize distant organs, the mechanisms by which primary tumor cells gain metastatic competence remain poorly understood. Increasing evidence demonstrates that fusion of tumor (i.e., neoplastic) and immune (e.g., macrophages) cells generate a distinct population of tumor-immune hybrid cells with enhanced functional ability to migrate and disseminate into peripheral blood. Herein, our study investigates tumor-macrophage hybrid cells, an underexplored population of disseminated tumor cells, and their inherent heterogeneity and acquisition of molecular mechanisms underlying their dissemination as metastatic effectors in colorectal cancer (CRC). Through hybrid cell phenotyping utilizing integrative single-cell RNA sequencing (scRNA-seq), cyclic immunofluorescence (cyCIF) and functional assays with an *in vitro* model of CRC hybrid cells, we identify Runt-related transcription factor 1 (*Runx1)* as a central regulator of hybrid cell motility and invasion. *Runx1* depletion in hybrid cells suppressed functional protease expression, chemotactic activity and extracellular matrix (ECM) invasion. Furthermore, pharmacologic inhibition of RUNX1 in an *in vivo* model reduced hybrid tumor growth and dissemination into peripheral blood, key attributes of metastatic spread of disease. In patients with CRC, RUNX1^+^ hybrid cells were identified in both primary tumor and peripheral blood, where circulating hybrid cells (CHCs) exhibited enriched migratory and epithelial-to-mesenchymal transition (EMT) phenotypes. Taken together, these findings reveal a mechanistic role for RUNX1 in driving invasive behavior of tumor-immune hybrids and highlight disseminated CHCs as an under-recognized contributor to metastatic spread and a promising noninvasive biomarker for tumor progression.

## Introduction

Metastatic progression is a primary cause of cancer-related mortality, where disseminated tumor cells establish lesions at distant sites.^1^ Up to 25% of colorectal cancer (CRC) patients are diagnosed with metastatic disease at initial diagnosis, while an additional 20-30% of patients with early-stage CRC harbor undetectable micro-metastatic disease that results in recurrence after frontline therapy.^2,3^ Metastatic disease arises through coordinated intrinsic genetic and epigenetic alterations, together with extrinsic microenvironmental cues, that reprogram tumor cell states to enable invasion, dissemination, and colonization of distant organs.^4,5^ Clonal selection and expansion during this evolutionary process generate profound intratumoral heterogeneity, constraining durable therapeutic responses. Despite these advances, the molecular circuits that govern acquisition of metastatic competence and associated treatment resistance remain incompletely defined.

An emerging mechanism by which tumor cells acquire metastatic competence is fusion with immune cells, giving rise to tumor-immune hybrid cells (hybrid cells/hybrids).^6–20^ These hybrid cells integrate genotypic and phenotypic programs from both parental lineages, coupling immune cell-derived migratory and immune-evasive properties with tumor cell-intrinsic proliferative and tumor-initiating capabilities.^21,22^ Beyond additive trait inheritance, hybrid cells adopt expanded cellular identities, including cancer stem cell-like states, indicating that fusion drives profound cellular reprogramming.^8,9,12,23–44^ As a result, tumor-immune hybrid cells have an enhanced capacity to traverse the metastatic cascade and constitute a biologically distinct and understudied class of disseminated tumor cells with the potential to redefine paradigms of metastatic evolution.

Tumor-immune hybrid cells are detectable in murine cancer models and in patients across a broad spectrum of solid malignancies, where they are found in primary tumors, peripheral blood, and metastatic lesions.^45,46^ Notably, circulating hybrid cells (CHCs) are markedly more abundant in peripheral blood than conventionally defined circulating tumor cells (CTCs), which lack immune protein expression, with multiple studies reporting CHC numbers up to two orders of magnitude higher than CTCs.^45,47^ Widely used CTC detection platforms, including FDA-approved CellSearch, rely on negative selection for CD45 to exclude hematopoietic cells^48^. Consequently, hybrid cells that retain immune markers such as CD45 are systematically eliminated, rendering this abundant and biologically consequential cell population largely invisible to standard CTC workflows and underrepresented in clinical literature.

Clinically, CHC abundance correlates with disease burden and overall survival in pancreatic, gastrointestinal, uveal melanoma, and head and neck cancers, underscoring their potential as a noninvasive biomarker of disease progression.^20,46,47,49–52^ Despite their prevalence and prognostic relevance, the genomic landscape, phenotypic diversity, and molecular programs that confer metastatic competence to hybrid cells remain largely undefined. Addressing this knowledge gap is critical to understanding the role of hybrid cells in tumor dissemination, therapeutic resistance, and disease recurrence. As the predominant tumor-derived cell population in circulation, defining the molecular drivers of hybrid cell behavior has substantial implications for improving early cancer detection and for developing strategies to intercept or limit metastatic disease.

In this study, we sought to define the transcriptomic, epigenetic, and functional heterogeneity of tumor-immune hybrid cells in CRC and to identify molecular mechanisms that govern their dissemination. We quantified hybrid cell phenotypes using single-cell RNA sequencing (scRNA-seq) analyses of public CRC datasets alongside newly generated datasets from fluorescence-activated cell sorting (FACS)-isolated hybrid cells from patient-matched primary CRC tumors and peripheral blood. To validate transcriptional programs at the protein level, we employed highly multiplexed whole-slide cyclic immunofluorescence (cyCIF) in matched primary tumor and peripheral blood specimens.

Our analyses reveal that hybrid cell states evolve dynamically across the metastatic cascade. Tumor-resident hybrid cells are enriched for migratory and invasive programs, whereas CHCs preferentially express immune-related and immune-evasive transcriptional signatures. Leveraging an *in vitro* CRC-macrophage cell fusion model, we demonstrate that hybrid cells acquire heightened transcriptional plasticity, a defining feature of metastasis.^53^ Notably, hybrid cells exhibit gene expression profiles that are distinct from both parental tumor cells and macrophages, as well as from tumor-macrophage cell doublets, confirming their unique cellular identity. Single cell-derived hybrid clones spanned a continuum from tumor-like to macrophage-like states at the transcriptomic, proteomic, and functional levels.

Among these states, hybrid cell clones with enhanced migratory and invasive capacity exhibited increased Runt-related transcription factor (*Runx1*) expression at both the transcript and protein levels, accompanied by elevated chromatin accessibility at the *Runx1* locus. Genetic depletion of *Runx1* suppressed protease expression and significantly impaired hybrid cell migration and extracellular matrix invasion. Consistently, pharmacologic inhibition of RUNX1 in a xenograft model of hybrid cell-derived tumors reduced tumor growth and dissemination *in vivo*. Importantly, RUNX1^+^ tumor-immune hybrid cells were detected in human CRC primary tumors and peripheral blood, where they were associated with epithelial-to-mesenchymal transition (EMT) signatures and increased prevalence with advancing disease stage. Collectively, these findings identify RUNX1 as a central regulator of hybrid cell dissemination in CRC and highlight its potential as a therapeutic target to constrain metastatic progression. By defining the molecular drivers of hybrid cell invasiveness and resolving their functional heterogeneity, this work reveals a previously underrecognized and clinically relevant disseminated tumor cell population that bridges tumor and immune identities, thereby reshaping current paradigms of metastatic evolution.

## Results

### Tumor-macrophage hybrid cells reprogram transcriptional states to enable dissemination and immune adaptation during metastatic progression in CRC

Using our previously established analytical workflow,^54^ we interrogated a publicly available scRNA-seq dataset of human CRC^55^ to resolve the diversity and transcriptomic states of tumor-macrophage hybrid cells within primary tumors and lymph node metastases. We identified tumor-macrophage hybrid cells based on co-expression of monocyte/macrophage markers (*CD68, CD14*, *CD163* and/or *CD11b;* see Table 1 for gene names) together with epithelial cell markers *ECAD*, *EpCAM*, or pan-cytokeratin (defined as keratins 2-5, 7-8, 14-16, and 19); across a recently published dataset comprised of 189 samples from 63 patients (Figure 1A).^56^ Following stringent doublet discrimination and exclusion, we identified 5,592 tumor-macrophage hybrid cells, approximately 2% of all sequenced cells, distributed across both primary tumor and lymph node specimens (Figure 1B and Supplemental Figure 1).

**Figure 1:**
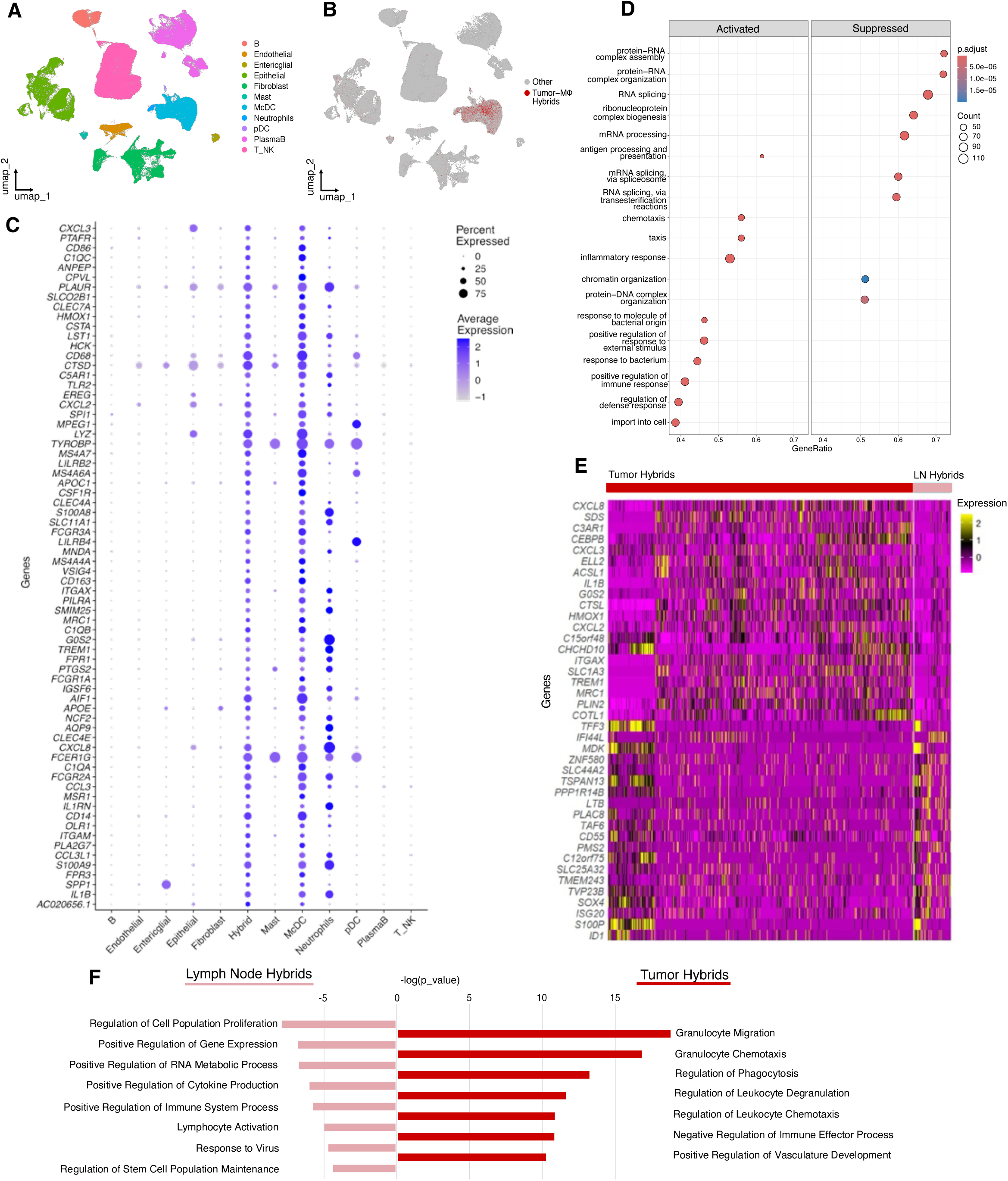
Tumor–macrophage hybrid cells upregulate immune evasive, migratory, and stem-like gene programs during colorectal cancer metastatic progression. A) UMAP depicting annotated cell type clusters of 373,058 cells from scRNA-sequencing of CRC tumors (n=63 patients).^55^ B) UMAP depicting hybrid cells (red) identified by co-expression of macrophage markers (CD45, CD163, or CD14) and one more epithelial markers (EPCAM, ECAD, panCK (KRT2–5, KRT7–8, KRT14–16, KRT19). C) Differential gene expression analysis of all identified hybrid cells vs all other sequenced cells. D) Gene set enrichment analysis (GSEA) Gene Ontology (GO) biological processes of hybrid cells using differential gene expression output. E) Differential gene expression analysis of tumor hybrid cells vs lymph node hybrid cells identified in patient matched tumor and lymph node specimens (n=7 patients). F) GSEA GO-biological processes of tumor hybrid cells vs lymph node hybrid cells using differentially expressed gene output.

**Table 1:** List of genes used throughout manuscript with gene symbol and full name.

| Gene ID | Gene Name |
| --- | --- |
| AIF1 | Allograft inflammatory factor 1 |
| AP-1 | Activator protein 1 |
| APC | Adenomatous polyposis coli |
| ARG2 | Arginase 2 |
| BRAF | B-Raf proto-oncogene |
| CD11b | Cluster of differentiation 11b |
| CD14 | Cluster of differentiation 14 |
| CD45 | Cluster of differentiation 45 |
| CD55 | Cluster of differentiation 55 |
| CD64 | Cluster of differentiation 64 |
| CD68 | Cluster of differentiation 68 |
| CDH1 | Cadherin 1 (E-cadherin) |
| CK | Cytokeratin |
| CSF1R | Colony-stimulating factor 1 receptor |
| CXCL2 | C-X-C motif chemokine ligand 2 |
| CXCL3 | C-X-C motif chemokine ligand 3 |
| CXCL8 | C-X-C motif chemokine ligand 8 |
| ECAD/CDH1 | E-cadherin |
| EpCAM | Epithelial cell adhesion molecule |
| ETS | E26 transformation-specific |
| FPR1 | Formyl peptide receptor 1 |
| FCGR3A | Fc gamma receptor IIIA |
| GAPDH | Glyceraldehyde-3-phosphate dehydrogenase |
| GRN | Granulin |
| IFI44L | Interferon-induced protein 44-like |
| ITGAM | Integrin subunit alpha M |
| KRAS | Kirsten rat sarcoma viral oncogene homolog |
| KRT | Keratin |
| LGALS3 | Galectin 3 |
| LGALS4 | Galectin 4 |
| LTB | Lymphotoxin beta |
| PECAM-1 | Platelet endothelial cell adhesion molecule 1 |
| PF4 | Platelet factor 4 |
| PLAUR | Plasminogen activator, urokinase receptor |
| PLEK | Pleckstrin |
| PTGS2 | Prostaglandin-endoperoxide synthase 2 |
| PTPRC | Protein tyrosine phosphatase receptor type C |
| RUNX1 | Runt-related transcription factor 1 |
| SOX4 | SRY-box transcription factor 4 |
| SPI1 | Spi-1 proto-oncogene |
| TGF-β | Transforming growth factor beta |
| THBS1 | Thrombospondin 1 |
| TP53 | Tumor protein p53 |
| TYROBP | TYRO protein tyrosine kinase binding protein |
| VIM | Vimentin |
| Wnt | Wingless/Integrated |

Differential gene expression (DEG) analyses revealed that hybrid cells exhibited elevated expression of genes associated with immune function and cellular migration, including *SPI1*, *PLAUR*, *GRN*, *AIF1* and *TYROBP,* relative to all other sequenced cell populations (Figure 1C). Notably, *AIF1* and *TYROBP* were also upregulated in hybrid cells identified in uveal melanoma primary tumors^54^, suggesting the existence of a conserved transcriptional program across tumor types. Consistent with these findings, global transcriptional profiling and UMAP visualization demonstrated that CRC hybrid cells more closely resembled macrophages than epithelial cells, while retaining distinct epithelial features (Figure 1B,C).

Gene set enrichment analysis (GSEA) using the Gene Ontology biological pathway annotations^57^ revealed significant enrichment of immune-related, chemotactic, and inflammatory pathways in hybrid cells (Figure 1D), consistent with prior observations in hybrid cells from uveal melanoma and prostate cancer.^54,58,59^ To further substantiate hybrid cell identity beyond marker-based classification, we directly compared hybrid cells, intestinal epithelial cells, and monocyte/classical dendritic cell (McDC) populations. Relative to epithelial cells, hybrid cells expressed elevated levels of immune-related genes including *CD163, CD45, FPR1, CD64, FCGR3A, PLEK*, and complement components (*C1QA-C)*. Conversely, compared to the McDC population, hybrid cells upregulated epithelial and colorectal-enriched genes (*KRT8, KRT10, KRT19, KRT18, EpCAM, TFF3, LGALS4, ARG2*), supporting a bona fide dual tumor-macrophage identity (full DEG lists in Supplemental File 1).

A unique feature of this publicly available dataset was the inclusion of seven matched primary tumor and lymph node samples, enabling evaluation of hybrid cell transcriptional states across the metastatic cascade. Within this subset, we identified 784 hybrid cells (0.85% of sequenced cells), comprising 700 tumor-resident hybrids (TrHCs) and 84 lymph node-resident hybrids. DEG analysis revealed that TrHCs preferentially upregulated *CXCL2, CXCL3, CXCL8* and *IL1B,* chemokines implicated in neutrophil recruitment, inflammatory signaling, and pro-metastatic niche formation (Figure 1E).^60,61^ In contrast, lymph node-resident hybrids exhibited increased expression of *IFI44L*, an interferon-stimulated gene^62^ and *LTB*, a gene enriched in lymphoid tissues.^63^ Notably, we identified a subset of TrHCs whose transcriptional profiles closely resembled those of lymph node-resident hybrids (Figure 1E, leftmost cluster). These cells expressed *SOX4*, a transcription factor associated with stemness, and *CD55*, an immune-evasive marker enriched in cancer stem cell populations^64^, suggesting that stem-like hybrid cells within the primary tumor may be primed for dissemination.

Consistent with these observations, GSEA revealed divergence across hybrid cell states: tumor-resident hybrids were enriched for pathways related to vasculature development, chemotaxis and cell migration, whereas lymph node-resident hybrids preferentially activated immune signaling and stem cell-associated pathways (Figure 1F). Collectively, these findings indicate that tumor-macrophage hybrid cells dynamically reprogram their transcriptional states during metastatic progression, with primary tumor-resident hybrids poised to facilitate dissemination through immune microenvironmental and vasculature remodeling, and lymph node-resident hybrids adopting phenotypes associated with immune adaptation and metastatic persistence.

### Tumor-macrophage cell fusion generates hybrid clones with heterogenous phenotypic states and metastatic properties

To interrogate the impact of tumor-macrophage fusion on cell phenotype and metastatic potential, we leveraged our previously established *in vitro* model of spontaneous fusion between the MC38-H2B-RFP CRC cell line and primary bone marrow-derived macrophages from β-actin-GFP C57BL/6 mice (Supplemental Figure 2A).^21,49^ After four days of co-culture, double-positive GFP^+^/RFP^+^ hybrid cells were FACS-isolated, expanded *in vitro*, then FACS-isolated into single cell clones (Figure 2A). Twelve hybrid clones were selected for downstream evaluation based on differences in morphology and GFP/RFP expression levels (Figure 2B and Supplemental Figure 2). Gene expression analysis using qRT-PCR revealed substantial heterogeneity in the expression of epithelial, immune, and EMT-associated genes across hybrid cell clones (Figure 2C). Macrophage-associated genes (*CD45, CD68, Csf1r, Ccl3*) varied widely, whereas most clones retained elevated epithelial gene expression (*EpCAM* and *Ecad)* despite their absence in the parental tumor cells.^65^ Hierarchical clustering delineated a continuum of hybrid phenotypes, spanning macrophage-like clones (e.g., Hybrid (H)4, H8, H11 and H12) and tumor-like clones (e.g., H1, H7, H9, H10, Figure 2C).

**Figure 2:**
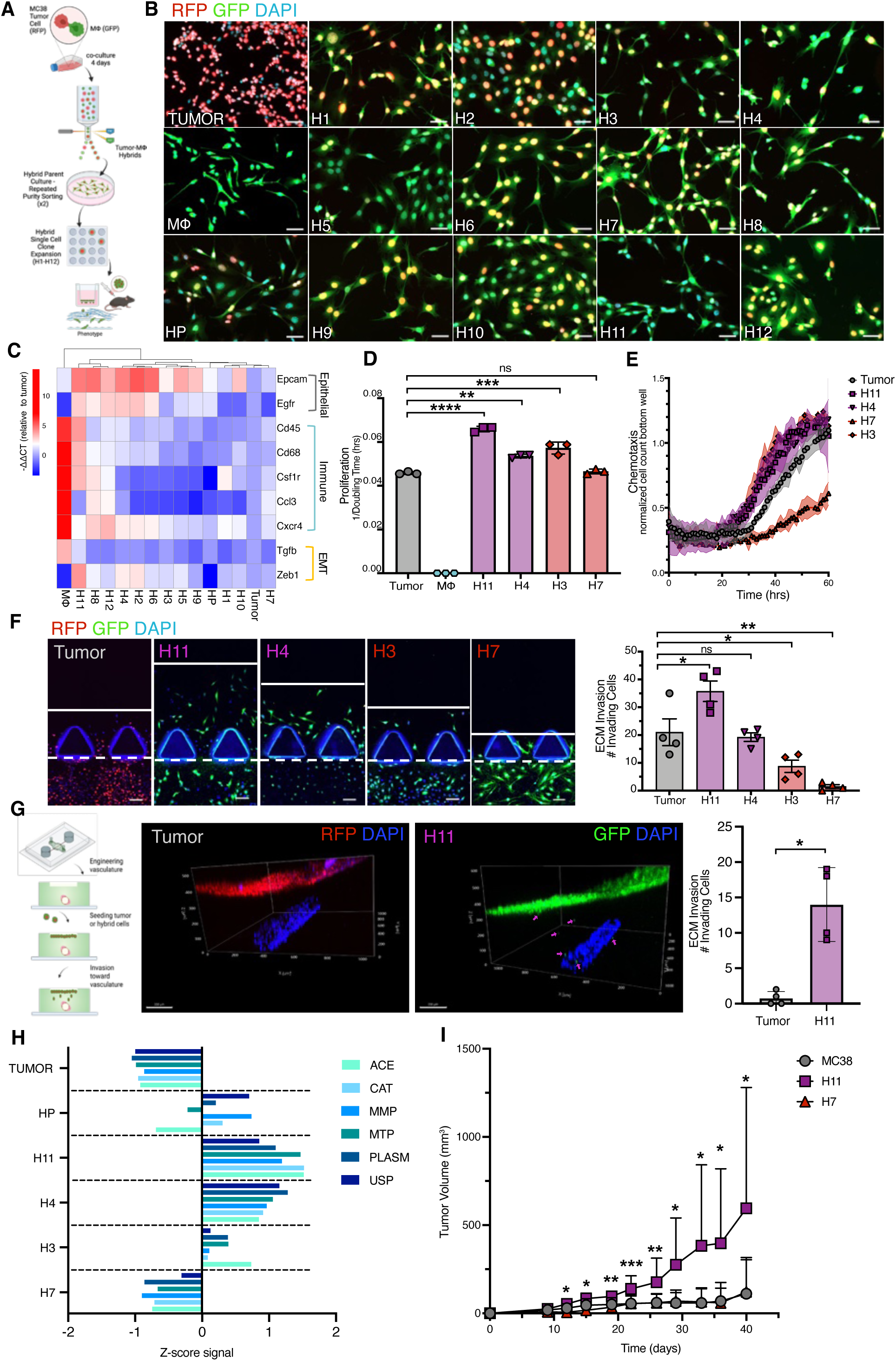
Colorectal cancer hybrid cell clones exhibit phenotypic and functional heterogeneity. A) Schematic of experimental design. B) Images of parental MC38-RFP tumor cells, parental Actin-GFP bone marrow derived macrophages (MΦ), MC38xMΦ hybrid parent (HP) cells, and single cell clones (H1-H12) generated from MC38xMΦ HP cell population. (scale bar = 50μM) C) Hierarchical clustering of macrophage, epithelial, and EMT gene expression from hybrid single cell clones and parental tumor and MΦ (n=3 biological replicates, each with 3 technical replicates, averaged). D) Proliferation of hybrid single cell clones (n=3 biological replicates, each with 3 technical replicates, averaged; data for all hybrid single cell clones in Supplemental Figure 2). E) Trans-well chemotaxis of hybrid single cell clones and MC38 tumor parent (n=3 biological replicates, each with 3 technical replicates, averaged; data for all hybrid single cell clones in supplemental figure 2). F) Hybrid cell invasion into collagen extracellular matrix (ECM) (imaging shown represents one section of whole well imaging from one of n=2 biological replicates, each with 4 technical replicates, scale bar = 100μm) and quantification of hybrid cell invasion in ECM represented as total number of invading cells after 72hrs. G) Invasion of MC38 tumor cells and H11 cells through collagen matrix toward vasculature, where pink arrowheads label migrating cells (imaging shown represents one of n=2 biological replicates, each with 4 technical replicates, scale bar = 200μm) and quantification of the number of invading cells. H) Protease activity in hybrid single cell clones and MC38 tumor parent (data shown represent one of n=2 biological replicates, each with 3 technical replicates). I) H11, H7 and MC38 *in vivo* murine tumor volume (n=27 MC38 (16 M, 11 F), n=15 H11 (10 M, 6 F) and n=8 H7 (4 M, 4 F)) data shown represent one of n=3 biological replicates) (*p* = **** <0.0001, *** <0.001, **<0.01, *<0.05)

Functional assays mirrored the transcriptional diversity observed across clones. When comparing hybrid clones, we noted marked heterogeneity in proliferation and chemotactic responses (Figure 2D and E, Supplemental Figure 2D, 2F, p-values vs. parent tumor cells: H11 *p* <0.0001, H4 *p* = 0.0002, H3 *p* = 0.0012, H7 *p* = 0.374). Specifically, macrophage-like clones (H11, H4) displayed enhanced invasive and migratory capacities relative to the tumor-like hybrid clones (H3, H7) and the parental tumor MC38 cells. Notably, H11 demonstrated significantly greater matrix degradation and ECM invasion, both in depth and cell number, when compared with H3, H7, and the parental tumor cells (Figure 2F, H11 *p* = 0.0112, H4 *p* = 0.9806, H3 *p* = 0.0359, H7 *p* = 0.0012), and showed more directional migration towards vasculature than tumor cells (Figure 2G, *p* = 0.013). Consistent with their invasive properties, protease expression directly correlated with increased ECM invasion across all hybrid clones, reinforcing a mechanistic link between transcriptional programs and invasive behavior (Figure 2H).

*In vivo,* we compared H11, the most macrophage-like clone, to the parental tumor MC38 line and the tumor-like H7 clone. H11 demonstrated heightened tumorigenic potential in C57Bl/6 mice, forming more rapidly growing tumors than both MC38 and H7 (Figure 2I, H11 vs MC38 and H7 *p* <0.05 days 12-40, * *p* <0.05, ** *p* <0.01, *** *p* <0.001, **** *p* <0.0001). Collectively, these data establish that tumor-macrophage fusion generates hybrid subpopulations with diverse gene expression programs, functional invasive capacities, and tumorigenic potential, highlighting hybrid cells as active effectors of metastatic progression.

### Tumor-macrophage cell fusion induces distinct transcriptomic and epigenetic landscapes underlying hybrid cell heterogeneity

To define transcriptomic and epigenetic landscapes of tumor-macrophage hybrid cells at single-cell resolution, we performed scRNA-seq and single-cell ATAC-seq (assay for transposase accessible chromatin sequencing) on four populations: 1) nascent hybrid cells (FACS-isolated after 4 days of co-culture), 2) established hybrid cells (maintained in culture through repeated purity FACS-isolation and passaging), 3) parental MC38 tumor cells, and 4) the parental macrophages (Figure 3A).

**Figure 3:**
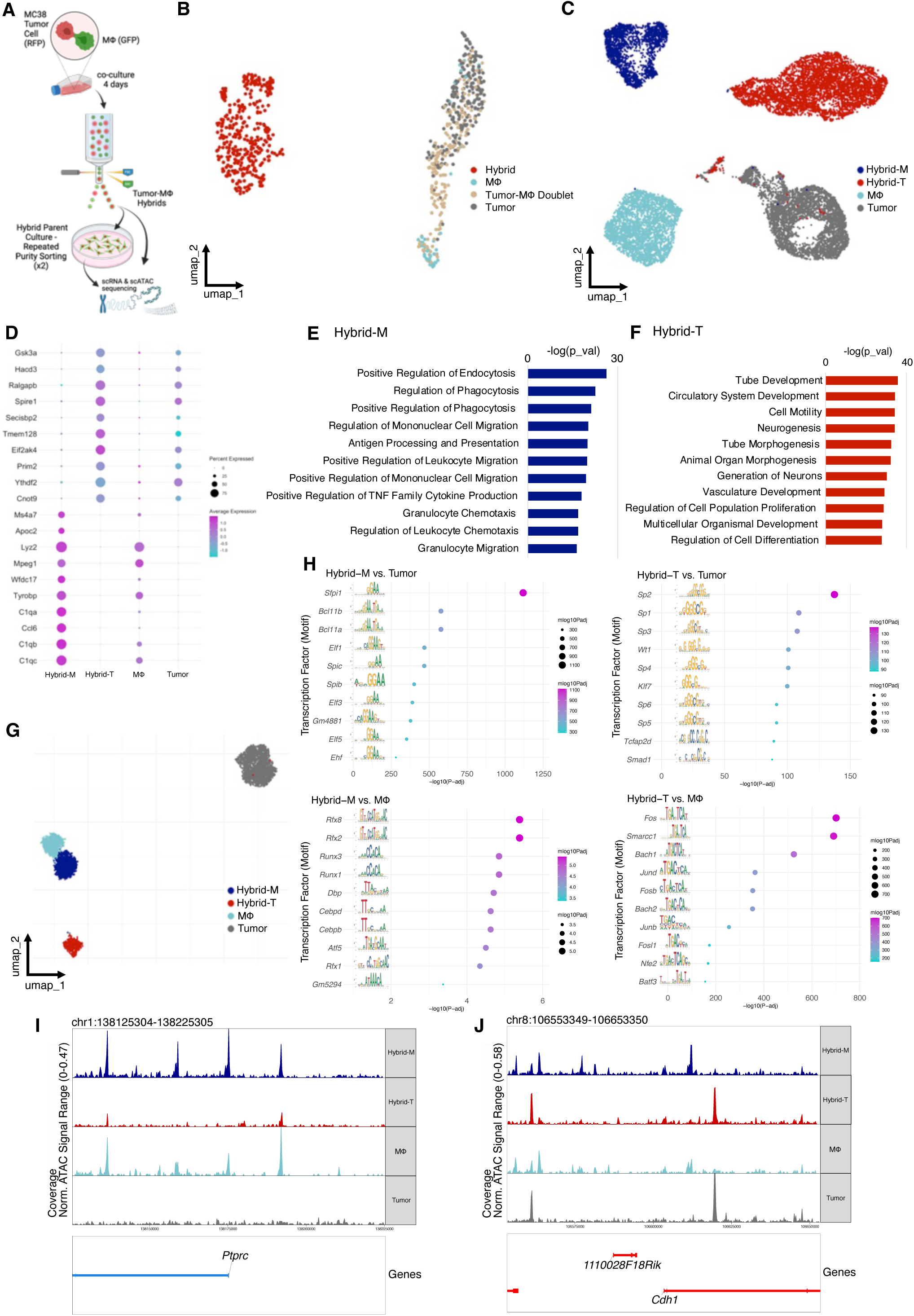
Multi-omic profiling reveals transcriptional and epigenetic heterogeneity of CRC hybrid cell lines. A) Schematic of experimental design. B) UMAP of scRNA-sequencing using Takara iCell8 for annotation of tumor-macrophage doublets, hybrid cells and tumor and macrophage cells (MΦ). C) UMAP of scRNA sequencing of “nascent” hybrids (hybrid-M), “established” hybrids (hybrid-T), MΦ, and MC38 tumor cells (data shown represent one of n=2 biological replicate sequencing experiments). D) Top 10 differentially expressed genes (DEGs) in hybrid-M and hybrid-T cells. E,F) GSEA GO-biological processes for hybrid-M and hybrid-T cells. G UMAP of scATAC-seq of “nascent” hybrids (hybrid-M), “established” hybrids (hybrid-T), MΦ, and MC38 tumor cells. H) scATAC-seq motif enrichment analysis for hybrid-M vs MΦ, hybrid-M vs tumor, hybrid-T vs MΦ, and hybrid-T vs tumor. I) scATAC-seq track plots for *Cd45 (Ptprc)* and J) *Ecad (Cdh1)* in hybrid-M, hybrid-T, tumor and MΦ cells. Additional scATAC-seq track plots included in Supplemental Figure 3.

A critical initial study distinguished bona fide hybrid cells from macrophage-tumor doublets, which can arise as artifacts in droplet-based library platforms. To address this, we employed the ICELL8 platform system (Takara), enabling multiple rounds of single cell dispensing into individual nanowells with imaging after each round to visually confirm wells containing single cells or artificial doublets (tumor + macrophage). Following Smart-Seq-based scRNA-seq with nanowell-specific barcoding, UMAP analysis demonstrated that fusion-derived hybrids formed distinct clusters, clearly separable from parental cells and artificially generated doublets (Figure 3B), validating the integrity of our single cell ∼omics pipeline and confirming that hybrid cell signatures are not artifacts of unresolved doublets.

High-throughput scRNA-seq using the 10X Genomics platform confirmed that hybrid populations were transcriptionally distinct from both the parental lineages (Figure 3C). Within the hybrid cell populations, nascent hybrid cells (hybrid-M) exhibited more immune-like transcriptional programs, aligning closely with macrophages, whereas established hybrids (hybrid-T) adopted a more tumor-like gene expression signature (Figure 3D). This shift likely reflects selection for proliferative hybrid clones during extended culture but may also represent biologically distinct subpopulations with clinical relevance. Notably, clonal heterogeneity persisted within established hybrids: for example, the H11 clone retained strong macrophage-like gene expression and functional traits, supporting a continuum of phenotypes spanning immune-like to tumor-like states.

GSEA of GO biological processes revealed that both hybrid-M and hybrid-T populations were enriched for pathways involved in cell motility. Hybrid-M cells additionally activated programs involved in phagocytosis, antigen processing and presentation, and development, whereas the hybrid-T cells were enriched for pathways related to proliferation, differentiation and developmental processes (Figure 3E, F). Collectively, these data underscore the dynamic and evolving nature of hybrid cell states, which occupy a phenotypic spectrum bridging immune and tumor identities. This flexibility likely underpins their migratory, immune-evasive, and metastatic potential, positioning them as important mediators of cancer progression.

Parallel scATAC-seq analyses of the same populations revealed chromatin accessibility landscapes that mirrored transcriptomic states. Both hybrid-M and hybrid-T populations displayed epigenetic profiles distinct from parental tumor and macrophage cells, with hybrid-M clustering closer to macrophages and hybrid-T clustering closer to tumor cells (Figure 3G). Track plot analyses of key loci, including CD45 (*Ptprc*) and Ecad (*Cdh1*), demonstrated coordinated chromatin accessibility changes at promoters, transcription start sites, and enhancers, consistent with transcriptional reprogramming (Figure 3I-J; Supplemental Figure 3).

Motif enrichment analysis revealed population-specific regulatory signatures. Hybrid-M cells were enriched for E26-transformation-specific (ETS) family transcription factors (Spi1 (*Sfpi1*) and *Spib*), and Runx family transcription factors (*Runx1* and *Runx3*), key regulators of immune function and cancer progression.^66,67^ In contrast, hybrid-T cells exhibited enrichment for Sp1-6 and AP-1 family motifs (*Fos*, *Fosb, Fosl1, Jund, and Junb*), central to tumor cell proliferation, invasion and metastasis (Figure 3H).^68,69^ Integration of scRNA-seq data confirmed that differential chromatin accessibility corresponded to transcriptional activity, and that hybrid cells gained not only elevated transcription factor expression but also an expanded repertoire of accessible target sites (Figure 3I-J and Supplemental Figure 3), establishing a distinct regulatory landscape divergent from either parental lineage following cell fusion. Collectively, these findings reveal that tumor-macrophage fusion induce profound transcriptomic and epigenetic heterogeneity, generating hybrid cell states that span a continuum between immune-like and tumor-like identities, with distinct regulatory programs that likely underpin their metastatic competence.

### *Runx1* drives hybrid cell migratory and invasive phenotypes

To identify key regulators underlying hybrid cell heterogeneity and invasive capacity, we performed an integrated analysis of transcriptomes, pathway enrichment, and chromatin accessibility. Across all modalities, RUNX1 consistently emerged as a top candidate, implicating its role as a central regulator of macrophage-like traits and metastatic potential in hybrid cells. Reactome pathway analysis^70^ revealed multiple RUNX1-associated pathways among the top 10 upregulated gene sets in hybrid cells (Figure 4A). Key upregulated genes within these pathways, including *Runx1, Thbs1, Spi1, Grn,* and *Pf4,* were markedly elevated in hybrid-M cells relative to hybrid-T cells, with Thbs1 upregulated across both hybrid populations (Figure 4B, Supplemental Figure 3). Concordantly, chromatin accessibility at the *Runx1*, *Spi1*, *Thbs1*, and *Pf4* loci was similarly increased in hybrid-M cells, indicating transcriptional activation of RUNX1-regulated programs (Figure 4C–D, Supplemental Figure 3). Given the established role of RUNX1 in EMT and CRC invasion,^71–73^ these findings suggest that acquisition of RUNX1 following cell fusion contributes to hybrid cell migration and invasion.

**Figure 4:**
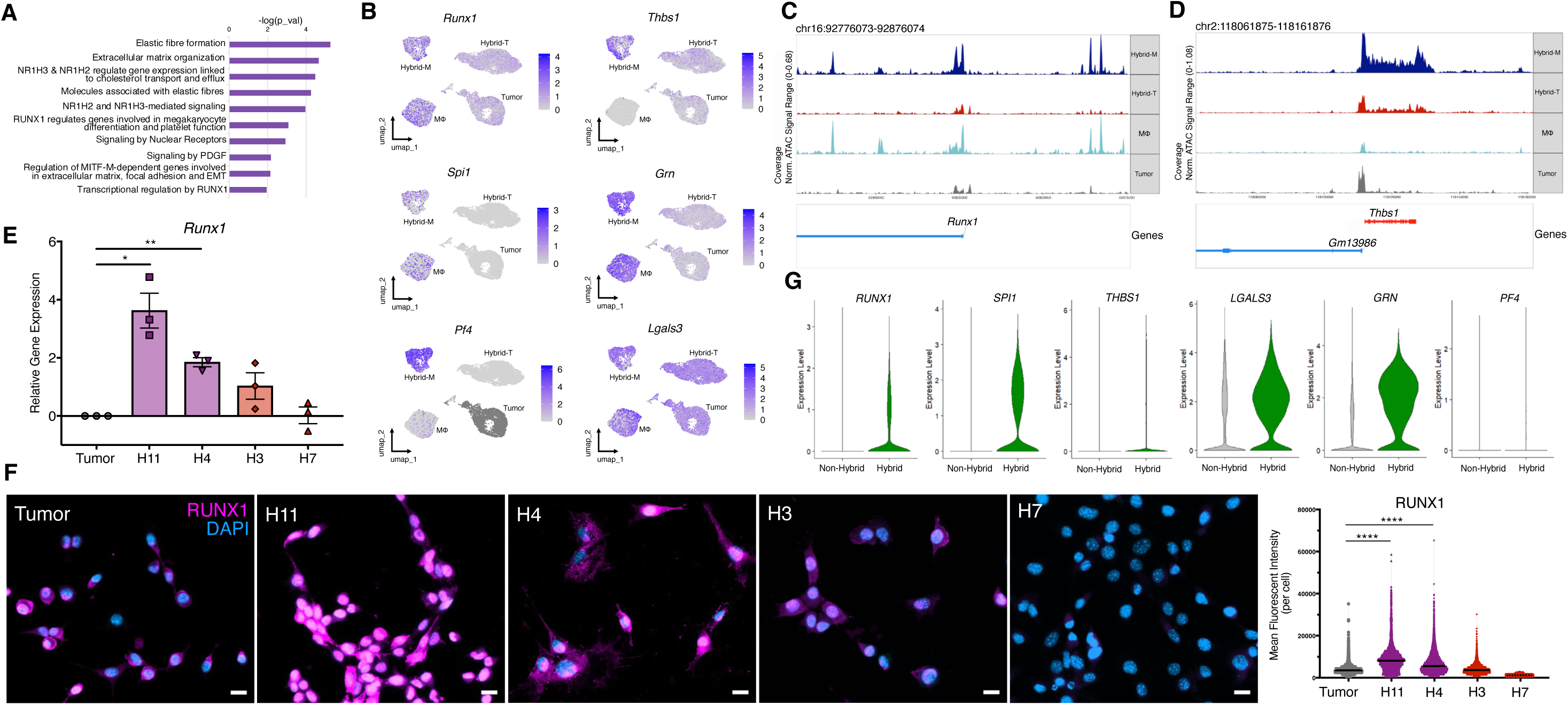
RUNX1 expression is elevated in invasive hybrid cell clones and associates with migratory and proteolytic programs. A) Reactome pathway analysis using DEGs for all hybrid cell (hybrid-T and hybrid-M) vs tumor cells (MC38) and macrophages (MΦ). B) Expression of key RUNX1 pathway associated genes contributing to Reactome analysis (*Runx1, Thbs1, Spi1, Grn, Pf4, Lgals3)*, overlaid onto the UMAP. C,D) scATAC-seq track plots for *Runx1* and *Thbs1* in hybrid-M, hybrid-T, tumor and MΦ cells. E) qRT-PCR *Runx1* gene expression in hybrid single cell clones H11, H4, H3, H7 relative to MC38 tumor cells (data shown represent n=3 biological replicates, each with 3 technical replicates averaged). F) Immunofluorescent staining of RUNX1 in hybrid single cell clones H11, H4, H3, H7 and MC38 tumor cells (scale bar = 50μm) and quantified as the mean fluorescent intensity per cell (images shown represent one area of whole well imaging from n=2 biological replicates). G) Gene expression of *RUNX1, THBS1, SPI1, GRN, PF4,* and *LGALS3* in human hybrid cells compared to all other cells from scRNA sequencing studies presented in Figure 1. All other RUNX1 pathway gene plots included in Supplemental Figure 3. (*p* = **** <0.0001, **<0.01, *<0.05)

Analyses of single-cell hybrid clones further supported this hypothesis. Macrophage-like hybrid clones (H11 and H4) exhibited significantly higher RUNX1 expression at both the mRNA (H11 *p* = 0.0001; H4 *p* = 0.0168) and protein levels (both *p* < 0.0001) relative to tumor-like clones (H3 *p* = 0.2101, H7 *p* >0.999) and the parental MC38 cells (Figure 4E,F; Supplemental Figure 3C). Functional assays demonstrated that increased RUNX1 expression directly correlated with enhanced invasive capacity, confirming its role as a phenotype determinant of hybrid cell aggressiveness.

To assess clinical relevance, we interrogated RUNX1 expression in hybrid cells from human CRC primary tumors and lymph nodes identified in Figure 1, a previously published dataset.^56^ Consistent with our *in vitro* model, human hybrid cells exhibited elevated *RUNX1* expression and upregulation of associated pathway members *THBS1, SPI1*, and *GRN* (Figure 4G, Supplemental Figure 3). Together, these data establish RUNX1 as a central regulator of the invasive and immune-like features of tumor-macrophage hybrid cells and provide a mechanistic rationale for investigating RUNX1 modulation as a strategy to limit hybrid cell-mediated migratory and invasive contribution to metastatic progression.

### Depletion of Runx1 impairs hybrid cell migration, invasion, and dissemination

To directly assess the functional contribution of *Runx1* to hybrid cell behavior, we generated knockdown cell lines of the macrophage-like hybrid clone H11, the tumor-like hybrid clone H7, and the parental MC38 tumor line using two independent lentiviral shRNA constructs targeting *Runx1.* Knockdown efficiency was confirmed at both the transcript and protein levels across all lines (Figure 5A–E, MC38 *p* <0.0001, H11 *p* <0.0001, H7 *p* <0.0001). Subsequent functional analyses were performed using the shRNA construct (shRNA-1) that achieved the most consistent suppression across all three cell lines.

**Figure 5:**
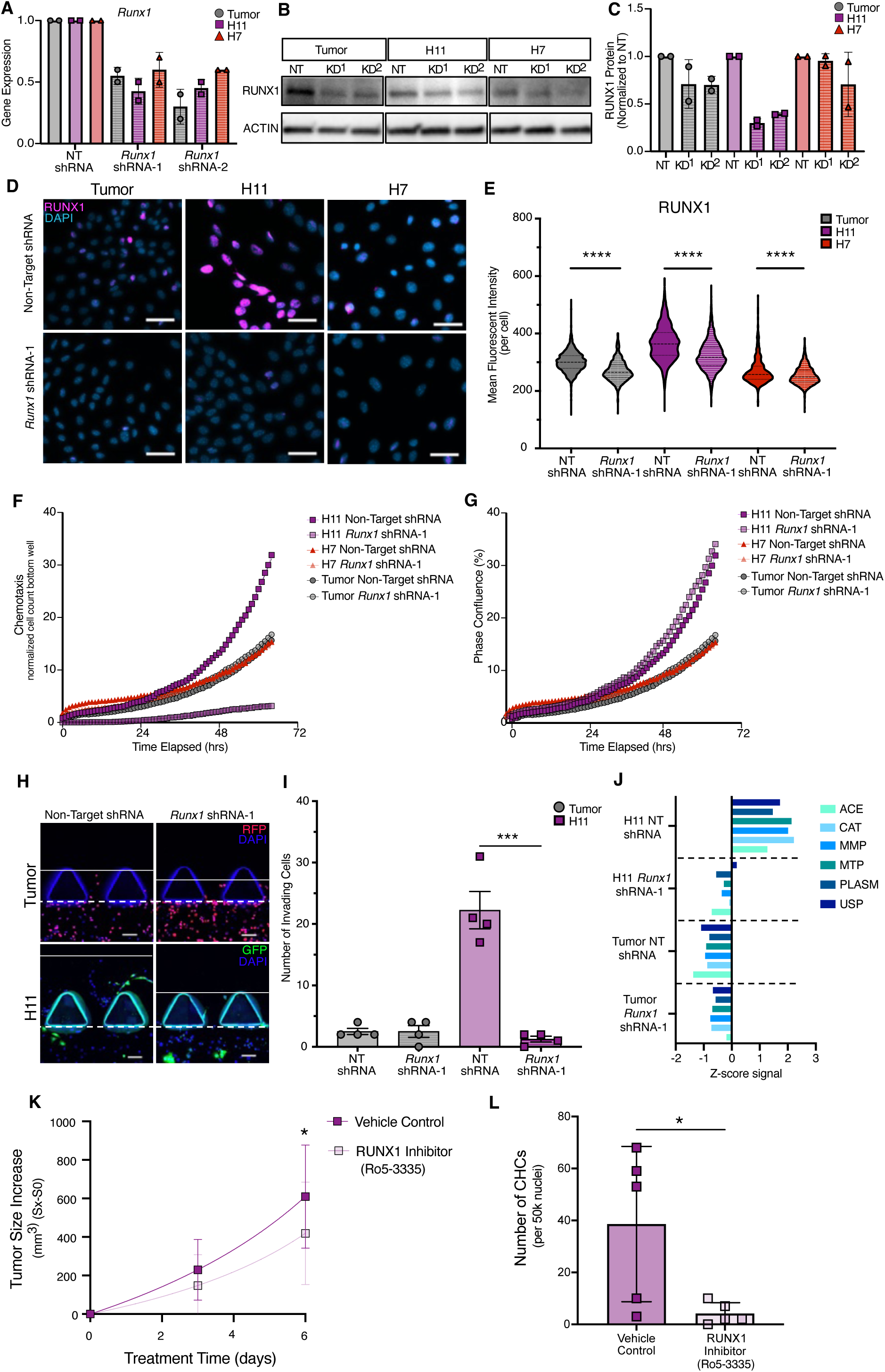
RUNX1 depletion impairs colorectal cancer hybrid cell invasion and limits tumor progression *in vivo.* A) qRT-PCR analysis of *Runx1* expression in MC38 tumor cells and hybrid cell lines H11 and H7 using two independent shRNA constructs and non-target (NT) control (data shown are an average of n=3 within-plate replicates and are representative of n=2 biological replicates each with n=2 technical replicates). B) Western blot of RUNX1 protein levels in knockdown (KD) samples across tumor, H11, and H7 cell lines, with ACTIN used as a loading control. C) Quantification of RUNX1 protein levels from western blot (B), each cell line normalized to each NT control after normalizing to ACTIN loading control (data shown representative of n=2 biological replicates each with n=2 technical replicates) D) Representative immunofluorescence images of RUNX1 staining in MC38, H11, and H7 NT-shRNA and *Runx1* shRNA-1 cells (scale bar = 50μm). E) Quantification of mean fluorescent intensity of RUNX1 per cell. *Runx1* shRNA-1 groups compared to non-target controls. F) Trans-well chemotaxis assay of MC38 tumor, H11 and H7 NT-shRNA and *Runx1* shRNA-1 cells. G) Proliferation assay of MC38 tumor, H11 and H7 NT-shRNA and *Runx1* shRNA-1 cells. H) Representative images from extracellular matrix (ECM) invasion assay for MC38 tumor and H11 NT-shRNA and *Runx1* shRNA-1 cells (scale bar = 100μm). I) Quantification of ECM-invading cells from (H) (data shown are representative of n=2 biological replicates each with n=2 technical replicates). J) Z-score normalized protease activity in MC38 tumor and H11 NT-shRNA and *Runx1* shRNA-1 cells (data shown are representative of n=2 biological replicates, each with n=3 technical replicates). K) *in-vivo* tumor growth of H11 with vehicle (n=10, 5M/5F) or Runx1 inhibitor (Ro5-3335) (n=10, 5M/5F) treatment (data shown are representative of n=2 biological replicates) L) The number of circulating hybrid cells (CHCs) (normalized to 50,000 nuclei) identified in the peripheral blood of mice bearing H11 tumors in RUNX1 treated (n=5) and vehicle (n=5) control groups. (*p =* **** <0.0001, *** <0.001, **<0.01, *<0.05)

In transwell chemotaxis assays, *Runx1* depletion profoundly impaired the migratory capacity of the highly invasive H11 hybrid cells, while exerting no measurable effect on the migration of the less invasive H7 hybrid or the MC38 tumor parent (Figure 5F). Notably, *Runx1* knockdown did not alter proliferation in any of the cell lines, indicating that its effects in H11 cells are selective for migratory behavior rather than general cell fitness (Figure 5G).

Consistent with these findings, ECM invasion assays revealed that *Runx1* depletion in H11 cells resulted in a marked reduction in both the number of invading cells and invasion depth (Figure 5H–I, H11 NT vs shRNA *p* = 0.0005), whereas invasion by MC38 cells was unaffected (*p* >0.9999). Loss of invasive capacity in H11 hybrids was accompanied by significant downregulation of invasion-associated proteases implicated in matrix degradation (Figure 5J), providing a mechanistic link between RUNX1 activity and hybrid cell invasiveness.

RUNX1 inhibition *in vivo* using Ro5-3335, a RUNX1-CBFβ interaction inhibitor that represses RUNX1/CBFB-dependent transactivation, resulted in both decreased H11 tumor size (Figure 5K, *p* = 0.03) as well as a decreased number of H11 hybrid cells detected in circulation (Figure 5L, *p* = 0.04, Supplemental Figure 4). Collectively, these data establish *Runx1* as a context-dependent but essential regulator of migration and invasion in macrophage-like hybrid cells and implicate RUNX1-driven programs as key effectors of hybrid cell-mediated tumor progression.

### RUNX1^+^ hybrid cells in matched CRC primary tumors and peripheral blood are associated with EMT phenotypes and disease progression

Having established RUNX1 as a central driver of hybrid cell invasiveness, we next evaluated the presence and phenotypic states of RUNX1^+^ hybrid cells in human CRC primary tumors and peripheral blood. We further examined whether RUNX1 and its downstream pathway components (SPI1, THBS1 or LGALS3) were associated with discrete hybrid phenotypic programs, including EMT, stem-like, proliferative, dormant, and immune-evasive states. To this end, we applied highly multiplexed cyCIF to patient matched primary tumor and peripheral blood specimens from 16 CRC patients (Table 2), profiling 34 markers to resolve hybrid cell identity and signaling states *in situ* (Table 3).

**Table 2:** Demographic information for cyclic & scRNA-seq patients.

| Patient ID | Sex | Age | Stage | Treatment<br>Prior to<br>Collection | Recurrence/<br>Progression/<br>Metastasis | Assays |
| --- | --- | --- | --- | --- | --- | --- |
| *CRC 94* | M | 68 | T3 N0 | Yes | No | cyCIF |
| *CRC 95* | F | 58 | T1 N1 | No | No | cyCIF |
| *CRC 96* | F | 60 | N/A | No | No | cyCIF |
| *CRC 97* | M | 65 | T3 N0 | Yes | Yes | cyCIF |
| *CRC 98* | F | 75 | T3 N0 | No | Yes | cyCIF |
| *CRC 99* | F | 73 | T1 NX | No | Yes | cyCIF |
| *CRC100* | F | 52 | T3 N0 | Yes | Yes | cyCIF |
| *CRC 101* | F | 69 | T4 N2A M1A | Yes | Yes | cyCIF |
| *CRC 102* | M | 53 | T2 N1A | No | No | cyCIF |
| *CRC 103* | M | 36 | T4b N0 | No | Yes | cyCIF |
| *CRC 104* | M | 68 | T3 N1a | Yes | Yes | cyCIF |
| *CRC 105* | M | 78 | T2 N0 | Yes | Yes | cyCIF |
| *CRC 106* | F | 53 | T3 N1c | No | No | cyCIF |
| *CRC 107* | F | 40 | T3 N1c | No | No | cyCIF |
| *CRC176* | F | 78 | T2 N0 | No | No | cyCIF, scRNA-seq |
| *CRC177* | M | 38 | T2 N1b | No | No | cyCIF, scRNA-seq |
| *CRC178* | M | 63 | T3 N0 | No | No | cyCIF |
| *CRC179* | F | 70 | T4a N1a | No | No | cyCIF, scRNA-seq |

**Table 3:** Ab-oligo antibodies for cyCIF.

| **Biomarker** | **Antibody Clone** | **Vendor** | **Catalog #** | **Species** |
| --- | --- | --- | --- | --- |
| αSMA | EPR5368 | Abcam | ab220795 | Rb |
| B2M | D8P1H | Cell Signaling | 12851 | Rb |
| CD20 | L26 | Abcam | ab9475 | Ms |
| CD31 | EPR3094 | Abcam | ab207090 | Rb |
| CD4 | EPR6855 | Abcam | ab181724 | Rb |
| CD44 | EPR1013Y | Abcam | ab216647 | Rb |
| CD45 | HI30 | BioLegend | 304002 | Ms |
| CD47 | polyclonal | Invitrogen | PA5-80435 | Ms |
| CD68 | D4B9C | Cell Signaling | 76437S | Rb |
| CD74 | VIC-Y1 | eBioscience | 14-0747-82 | Ms |
| CD8 | C8/144B | Thermo Fisher | MA5-13473 | Ms |
| CDX1 | EPR4819(2) | Abcam | ab248164 | Rb |
| CDX2 | EPR2764Y | Abcam | ab220799 | Rb |
| ECAD | EP700Y | Abcam | ab201499 | Rb |
| EpCAM | VU1D9 | Cell Signaling | 46403SF | Ms |
| ERK1/2 | 137F5 | Cell Signaling | 68303 | Rb |
| FOXP3 | 236A/E7 | Thermo Fisher | 50-112-9542 | Ms |
| Ki-67 | pAb | Abcam | ab15580 | Rb |
| L1CAM | UJ127.11 | MilliporeSigma | L4543 | Ms |
| LGALS3 | EP2775Y | Abcam | ab197544 | Rb |
| LYVE1 | polyclonal | Abcam | ab14917 | Rb |
| Neurofilament (NFT) | polyclonal | MilliporeSigma | AB5539 | Chk |
| p16 (CDKN2A) | 4844R | Novus Biologicals | NBP3-14223 | Rb |
| p27 | KIP1 [Y236] | Abcam | ab206927 | Rb |
| p38 (MAPK) | Y122 | Abcam | ab227995 | Rb |
| panCK | AE1/AE3 | Santa Cruz | sc81714 | Ms |
| RUNX1 | D33G6 | Cell Signaling | 4336BF | Rb |
| S100 | EP1576Y | Abcam | ab215989 | Rb |
| SOX9 | D8G8H | Cell Signaling | 74021 | Rb |
| SPI1 | 9G7 | Cell Signaling | 2258 | Rb |
| TEAD4 | polyclonal | Invitrogen | PA5-21977 | Rb |
| TGF-β | polyclonal | R&D Systems | AB-100-NA | Rb |
| THBS1 | EPR22927-54 | Abcam | ab267397 | Rb |
| Vimentin (VIM) | D21H3 | Cell Signaling | 5751BF | Rb |

Across all patient peripheral blood samples, CHCs were significantly more abundant than conventionally defined CTCs at every disease stage, consistent with prior studies^46,74^ (Figure 6A, stage 1 CHC vs CTC *p* <0.0001, stage 2 CHC vs CTC *p* <0.0001, stage 3 CHC vs CTC *p* <0.0001). CHC abundance increased with disease progression, although this relationship was partially confounded by treatment status at the time of sample collection. Unsupervised phenotypic clustering of CHCs and CTCs revealed substantial heterogeneity within the CHC population, resolving multiple distinct cell states based on marker expression (Figure 6B-C, E and Supplemental Figure 5). Notably, RUNX1^+^ hybrid cells were identified in primary tumors and peripheral blood (Figure 6D). Seven phenotypic clusters (clusters 0, 3, 4, 6, 7, 9 and 13) were characterized by high expression of RUNX1 and/or downstream effectors SPI1 and THBS1 (Figure 6E-F), defining a recurrent RUNX1-activated hybrid cell state. Five of these RUNX1^+^ clusters (clusters 0, 3, 6, 7 and 12) additionally co-expressed vimentin (VIM), a canonical EMT marker, and were composed predominantly of CHCs and not CTCs (which were enriched in cluster 4). These findings suggest that RUNX1^+^ hybrid cells in human CRC preferentially adopt EMT-like phenotypes, mirroring the invasive programs observed in murine *in vitro* fusion-derived hybrids. Consistent with a role in disease progression, expression of RUNX1 pathway components (SPI1 and THBS1) within CHCs increased with advancing CRC stage (Figure 6G, SPI1 stage1v2 *p* = 0.03, SPI1 stage1v3 *p* = 0.03, THBS1 stage1v2 *p* = 0.05, THBS1 stage1v3 *p* = 0.05). Together, these data identify RUNX1^+^ hybrid cells as a transcriptionally and phenotypically distinct subpopulation enriched in the circulation of CRC patients, particularly at later disease stages. Collectively, the abundance, EMT-associated features and progressive enrichment of key metastatic pathways underscore that RUNX1^+^ hybrid cells may have potential clinical relevance as biomarkers of metastatic competence and disease progression, and implicate this hybrid cell state as a promising target for disease monitoring and therapeutic intervention in CRC.

**Figure 6:**
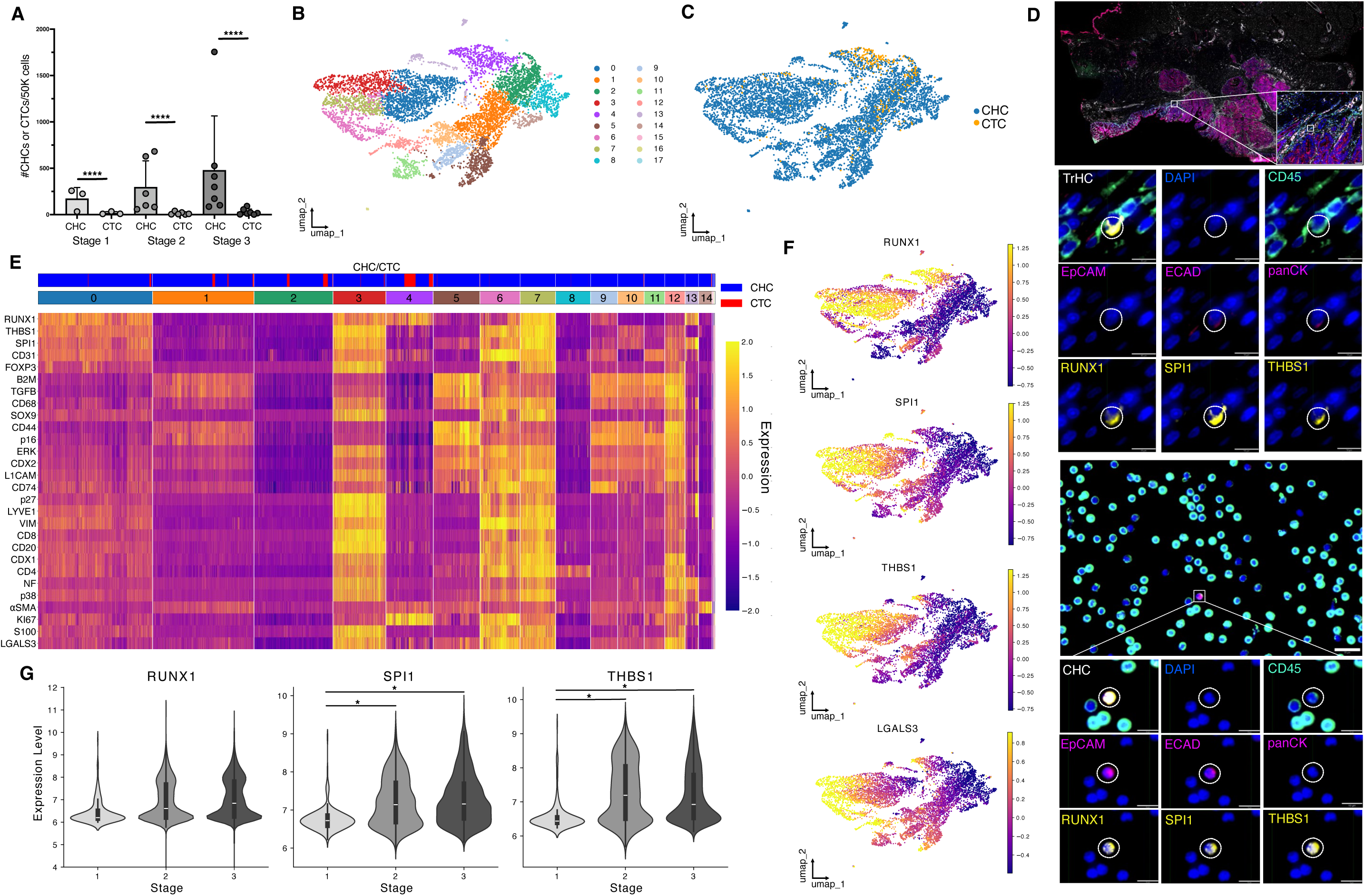
Cyclic immunofluorescence identifies RUNX1⁺ tumor resident and circulating hybrid cells associated with colorectal cancer progression. A) Quantification of circulating hybrid cells (CHCs) and circulating tumor cells (CTCs) per 50,000 nucleated cells in peripheral blood across CRC stages 1, 2, and 3. B) UMAP plot of cyclic immunofluorescence (cyCIF) data colored by phenotypically defined clusters. C) UMAP plot from (B) labeled by CHC versus CTC identity. D) Representative cycIF image of a whole-slide CRC primary tumor showing RUNX1+ tumor hybrid cells (TrHC) and Runx1+ CHC (hybrid cells identified as DAPI+, CD45+, EpCAM+ or ECAD+ or panCK+) including Runx1 downstream pathway members SPI1 and THBS1. (scale bars; tumor inset = 50μm, TrHC individual images = 10μm, peripheral blood = 20μm, CHC individual images = 10μm). E) Heatmap of 34 phenotypic markers measured by cyCIF across individual CHCs and CTCs, clustered by expression patterns. F) UMAP feature plots showing normalized average fluorescent intensity levels of RUNX1, SPI1, THBS1, and LGALS3 across all single cells. G) Violin plots of RUNX1, SPI1, and THBS1 expression in CHCs grouped by CRC disease stage (*p* = * < 0.05).

### Patient-derived tumor-immune hybrid cells exhibit functionally distinct immune-like states and conserved RUNX1-driven transcriptional programs in primary tumors and circulation

To define the transcriptional heterogeneity of patient-derived tumor-immune hybrid cells, we performed scRNA-seq on FACS-isolated hybrid cells (ECAD⁺/EpCAM⁺/CD45⁺) from matched primary CRC tumor (tumor-resident hybrid cells, TrHCs) and peripheral blood samples (CHCs) (Supplemental Figure 6). Given the rarity of hybrid cells in clinical specimens, we employed the SMART-Seq platform (Takara Bio), which enables the generation of full-length transcriptomes from picogram-scale RNA inputs. For each patient, control populations of tumor cells (ECAD⁺/EpCAM⁺) and immune cells (CD45^+^ PBMCs) were also profiled (n = 16 cells per population, per patient). Importantly, panCK was not used as a hybrid selection marker in this assay because its cytoplasmic localization necessitates fixation and permeabilization for FACS, which can degrade RNA and limit transcriptomic profiling.

Unsupervised clustering revealed multiple transcriptionally distinct CHC states (CHC1-3), each represented across all three patients analyzed (Figure 7A-C). Both TrHCs and CHCs (notably the CHC1, CHC2, CHC3 clusters) exhibited transcriptional similarity to CD45^+^ immune populations, clustering more closely with immune cell populations than with EpCAM^+^ tumor controls, with only a minor fraction of CHCs (CHC light pink cluster) aligning with epithelial tumor clusters. These data suggest that the majority of hybrid cells adopt immune-like transcriptional programs during dissemination, or that immune-associated states are selectively permissive for circulation. Analysis of hybrid cells in circulation (CHCs) compared to primary tumor hybrids (TrHCs) using differential gene expression and pathway enrichment analysis revealed that CHCs were enriched for chemotaxis, developmental programs, and ECM remodeling relative to TrHCs (Supplemental Figure 7A-B), highlighting both shared and context-dependent transcriptional features. Importantly, each hybrid subpopulation, both TrHCs and CHCs, were transcriptionally distinct from both tumor and immune controls (Figure 7D). Pathway analysis of the three distinct CHC clusters (CHC1-3) revealed functional specialization: CHC1 cells were enriched for macrophage-associated phagocytic, migratory, and cytokine response programs; CHC2 cells for B-cell activation pathways; and CHC3 cells for T-cell activation and adhesion processes (Figure 7E). These findings indicate that CHCs occupy discrete immune-like functional states.

**Figure 7:**
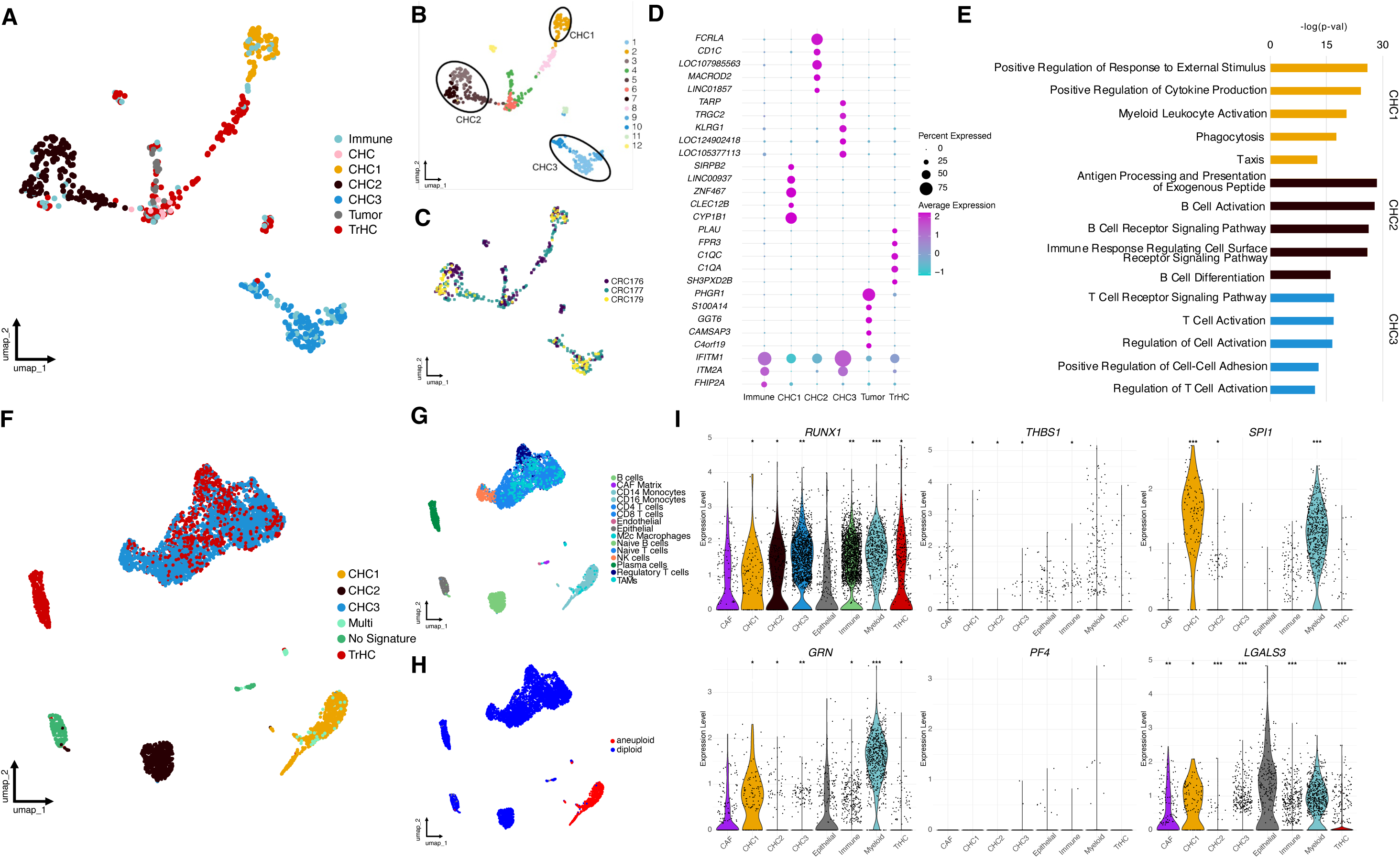
Single-cell RNA sequencing reveals functionally distinct tumor-immune hybrid cell populations with conserved RUNX1 transcriptional programs in tumor-macrophage hybrids. A) UMAP projection of SMART-Seq data showing clustering of tumor-resident hybrid cells (TrHCs, EpCAM⁺/ECAD⁺/CD45⁺), circulating hybrid cells (CHCs, EpCAM⁺/ECAD⁺/CD45⁺), tumor cells (EpCAM⁺/ECAD⁺), and immune cells (CD45⁺) from patient-matched colorectal tumor and peripheral blood samples (n = 3 patients; n=3 PBMC samples, n=2 tumor samples). B) UMAP colored by CHC subclusters (CHC1–3) with distinct transcriptional states. C) UMAP colored by patient ID. D) Dot plot showing the top five differentially expressed genes defining each major cell type. E) Gene set enrichment analysis of gene ontology - biological processes highlighting functional differences among CHC1–3 clusters. F) UMAP of 10X Genomics data enriched for tumor-resident hybrids and circulating hybrid cells, colored by hybrid signature scores derived from SMART-seq CHC cluster modules. G) UMAP of the 10X Genomics dataset annotated by major cell types, showing distribution of hybrid-enriched populations. H) Aneuploidy prediction in 10X Genomics dataset distinguishing diploid versus aneuploid populations. I) Violin plots of RUNX1 genes identified in tumor-macrophage murine hybrids (*RUNX1, THBS1, SPI1, GRN, PF4, LGALS3*). (*p* = *** <0.001, **<0.01, *<0.05)

In a subset of patients with sufficient hybrid cell yield, we additionally performed high-throughput 10X Genomics-based scRNA-seq on pooled non-hybrid primary tumor or peripheral blood spiked with an enrichment of hybrid cells (ECAD⁺/EpCAM⁺/CD45⁺) to capture the broader cellular landscape (Supplemental Figure 7D). Although these samples were enriched for TrHCs or CHCs, we expected them to also contain a mixture of immune, epithelial, stromal, and CRC tumor cells, particularly in tumor-derived biopsies. Because these cells were not sorted into uniquely barcoded wells, as in the SMART-Seq, surface marker-based annotation was not feasible. To overcome this limitation, we generated hybrid gene expression modules from the SMART-Seq CHC clusters and applied them to the 10X datasets to identify hybrid cells (Figure 7F). Additional cell types were annotated using established epithelial, immune, stromal, and macrophage gene modules (Figure 7G).

This integrative approach revealed multiple hybrid-enriched clusters within the 10X datasets, including a macrophage-like hybrid population and additional clusters aligned with B cell- and T cell-associated transcriptional programs (Figure 7F,G). Aneuploidy inference (Figure 7H) and tumor-specific gene expression scoring (Supplemental Figure 7E-F) demonstrated that macrophage-like hybrid cells exhibited elevated aneuploidy and higher expression of tumor-specific genes, whereas B cell- and T cell-like hybrid populations showed substantially lower aneuploidy rates, supporting distinct evolutionary origins or selective pressures among hybrid states. Importantly, the hybrid cells sequenced did not show an increase in doublet prediction scoring compared to the rest of the sequenced cells (Supplemental Figure 7G), which align with our earlier findings detailed in Figure 1 and Figure 3.

Across both scRNA-seq platforms, *RUNX1* and its downstream effectors (*SPI1, THBS1, GRN, LGALS3*) were significantly enriched in both CHCs and TrHCs relative to parental tumor control cells (Figure 7I and Supplemental Figure 7C). These *RUNX1*-associated transcriptional programs overlapped with pathways identified *in vitro*, including migration, chemotaxis, and immune activation. Notably, RUNX1 and downstream associated genes were most highly expressed in CHC1 cluster, the macrophage-like hybrid population with elevated aneuploidy prediction and higher tumor gene expression module scoring, implicating RUNX1-driven regulatory networks as conserved determinants of invasive hybrid phenotypes. Collectively, these findings demonstrate transcriptional convergence between patient-derived and experimentally generated tumor-immune hybrid cells and establish RUNX1-centered programs as central regulators of hybrid cell invasiveness and metastatic competence in CRC.

## Discussion

Fusion-derived hybrid cells have been proposed as a source of tumor plasticity, yet whether they represent a uniform population or functionally distinct states remains unknown. Here we show that tumor-macrophage hybrid cells comprise heterogenous and specialized cell states, with distinct subpopulations enriched for programs associated with metastatic progression. By integrating single cell transcriptomics and epigenomic profiling with functional interrogation of *in vitro*-derived hybrids and patient-matched primary tumor hybrids and peripheral blood CHCs, we uncover hybrid cell subpopulations enriched for migratory, immune-evasive, and stem-like programs that align with key steps of the metastatic cascade. These data reveal a previously underappreciated cellular mechanism of tumor cell dissemination from the primary tumor, in which fusion-derived hybrid cells bridge tumor and immune phenotypes to promote invasion and spread. A central finding of this work is the identification of RUNX1, a transcription factor classically associated with hematopoiesis and leukemogenesis, as a critical regulator of hybrid cell invasiveness and a potential driver of metastatic progression in CRC. RUNX1 expression was selectively enriched in macrophage-like hybrid cell subpopulations and was functionally required for migration, ECM degradation, and invasive behavior. Genetic depletion of *Runx1* in invasive hybrid cells (hybrid-M) markedly impaired chemotaxis and invasion, while downregulating protease activity, implicating RUNX1 as a nonredundant regulator of hybrid cell motility. Moreover, pharmacological inhibition of RUNX1 significantly suppressed tumor growth and dissemination *in vivo,* underscoring its therapeutic relevance. These findings are consistent with prior studies implicating RUNX1 in CRC progression, EMT, angiogenesis, and immune modulation.^75–80^ Previous work has shown that RUNX1 promotes metastasis through activation of Wnt/β-catenin signaling, induction of PTGS2 (COX2) expression, facilitation of vessel co-option, and modulation of tumor-stromal crosstalk^73,81^ via TGF-β and THBS1 signaling,^82^ with validation in both *in vitro* and *in vivo* CRC models.^71^ Our data extend these observations by positioning RUNX1 within a specific cellular context, tumor-macrophage hybrid cells, where its activation appears to preferentially endow hybrid cells with enhanced invasive and metastatic capacity. Together, these findings suggest that RUNX1 is selectively induced in macrophage-like hybrid cells in response to microenvironmental cues, conferring a unique convergence of immune adaptability and tumor aggressiveness.

Across both *in vitro*-derived hybrids and patient-derived specimens, hybrid cells occupied a phenotypic continuum between macrophage-like and tumor-like states and exhibited pronounced transcriptional plasticity. RUNX1 and its downstream effectors (SPI1 and THBS1), along with pathways involved in migration, ECM remodeling, and immune signaling, were consistently enriched in macrophage-like hybrid states, which displayed the highest invasive potential in both functional assays and GSEA. Single-cell profiling using both scRNA-sequencing and cyCIF identified RUNX1^+^ hybrid cells as a transcriptionally and phenotypically distinct subpopulation present in both primary tumors and circulation. Notably, RUNX1^+^ hybrid cells were enriched for EMT, stemness, and immune modulatory programs, indicating that RUNX1-associated transcriptional states are conserved features of invasive hybrid cells across disease compartments.

Importantly, multiple transcriptionally distinct hybrid cell clusters were identified across patients, many of which were enriched for RUNX1 or its downstream effectors. The prevalence of RUNX1^+^, SPI1^+^ and THBS1^+^ hybrid cells increased with advancing CRC stage, highlighting RUNX1 as a multifaceted driver of metastatic progression that integrates EMT, migration, angiogenesis, and tumor-immune hybrid cell dissemination in cancer and establishes tumor-immune hybrid cells as active effectors of CRC metastasis rather than passive byproducts of tumor evolution.

Beyond mechanistic insights, our findings have important translational implications. The consistent detection of RUNX1^+^ hybrid cells in circulation, and their increasing prevalence with disease stage, suggests that these cells may serve as a noninvasive biomarker of metastatic risk. Moreover, the functional requirement for RUNX1 in hybrid cell invasion identifies this pathway as a candidate therapeutic target. Encouraging pharmacologic inhibition of RUNX1 using Ro5-3335 has demonstrated efficacy in leukemia, PDAC, and glioblastoma,^83–85^ supporting the feasibility of targeting this pathway in other solid tumors. Consistent with this, we show that pharmacologic inhibition of RUNX1 using Ro5-3335 in H11 subcutaneous murine xenografts led to reduced tumor growth and a substantial decrease in the number of hybrid cells detected in circulation, supporting a role for RUNX1 in promoting hybrid cell dissemination *in vivo*. While further studies are needed to define the precise mechanisms underlying these effects, these data support RUNX1 as a promising therapeutic target in hybrid-cell associated metastasis.

Several study limitations should be considered. Hybrid cells in this study were identified by their co-expression of epithelial tumor and immune cell markers, which may underestimate the full spectrum of fusion-derived hybrids, particularly those that undergo phenotype switching or marker loss over time.^86^ More sophisticated lineage-tracing approaches, including MADM-CloneSeq^87,88^ or fusion reporter systems like Cre-Lox fate mapping^89^ using *in vivo* systems would improve specificity and temporal resolution of hybrid cell identification. Additionally, while RUNX1 was necessary for hybrid cell invasion, its sufficiency alone in driving macrophage-like hybrid phenotypes remains unclear. Future investigations should explore how RUNX1 integrates with other transcriptional regulators to control immune evasion, stemness, and survival in circulation, and how tumor microenvironmental cues shape RUNX1 activation. Finally, although we analyzed patient-matched tumor and peripheral blood samples, larger clinical cohorts will be required to fully assess interpatient heterogeneity and determine whether RUNX1^+^ CHCs reliably predict metastatic potential or correlate with clinical outcomes.

In summary, this study provides comprehensive single cell and functional dissection of tumor-immune hybrid cells in CRC and identifies RUNX1 as a pivotal regulator of hybrid cell invasion and metastatic competence. To our knowledge, this is the first analysis of scRNA-seq datasets of hybrid cells from patient-matched tumor and peripheral blood samples, providing a valuable foundation for exploration of hybrid cell biology across the metastatic cascade. By distinguishing bona fide fusion-derived hybrid cells from technical artifacts/doublets and linking their transcription programs to functional behavior, our work defines distinct hybrid cell states that align with metastatic progression. Beyond this technical advance, we establish tumor-immune fusion as a previously underappreciated driver of metastasis and position RUNX1 as a promising target for therapeutic intervention and biomarker of CRC progression.

## Data Availability

Code used for analysis is available at https://github.com/KoDeQuestion/CRCHybridCollab.

## Acknowledgments

We thank the Advanced Light Microscopy, Histopathology, and Flow Cytometry Cores at Oregon Health and Science University (OHSU) for technical support. We are grateful to our clinical collaborators at OHSU for assistance with patient specimen acquisition and to the patients who generously donated samples for this study. We also thank QiTissue for their support with cyCIF image registration and processing. We acknowledge Kathryn Fowler, Abigial Gillingham, Hannah Farley, Ethan Lu, and Ashvin Nair for their technical assistance with antibody validation and PBMC slide preparation. We further thank the Knight Biostatistics Shared Resource, Dr. Byung Park and Yun Yu, for their guidance and support with the statistical analyses. Figures were created using BioRender and Affinity Design 2.

## Author Contributions

Conceptualization, A.N.A., K.Q., A.C.A., S.L.G., M.H.W.

Methodology, A.N.A., K.Q., L.E.B., C.M.F., J.M.F., G.W., Y.H.C., A.C.A., S.L.G., M.H.W.,

Formal analysis, A.N.A., K.Q., N.G., J.A.J., C.C.R., A.G.M., D.R., S.G., I.D.M., G.W., C.M.F., J.M.F., A.C.A, S.L.G, M.H.W

Data curation, A.N.A., K.Q., N.G., J.A.J, A.P., A.Z., G.H., B.J.S., J.R.S., A.G.M., D.R., W.G., P.H.T., C.M.F., J.M.F.

Writing – original draft preparation, A.N.A., K.Q., S.L.G., M.H.W.

Writing – reviewing and editing, all authors.

Resources, K.T., G.W., L.E.B., V.L.T., B.B., C.D.L., C.M.F., J.M.F., Y.H.C., A.C.A., S.L.G., M.H.W.

Supervision and project administration, A.N.A., S.L.G., M.H.W.

Funding acquisition, A.N.A., K.Q., Y.H.C., A.C.A., S.L.G., M.H.W.

## Abbreviations

2D: Two-dimensional
3D: Three-dimensional
Ab-oligo: Antibody-oligonucleotide
AF555: Alexa Fluor 555
AF647: Alexa Fluor 647
AF750: Alexa Fluor 750
ANOVA: Analysis of variance
ATAC-seq: Assay for transposase-accessible chromatin sequencing
BCA: Bicinchoninic acid
BME: Beta-mercaptoethanol
BSA: Bovine serum albumin
cDNA: Complementary DNA
CHC: Circulating hybrid cell
CO2: Carbon dioxide
COX2: Cyclooxygenase 2
CRC: Colorectal cancer
CTC: Circulating tumor cell
cyCIF: Cyclic immunofluorescence
DAPI: 4′,6-diamidino-2-phenylindole
DEG: Differentially expressed gene
DIW: Deionized water
DMEM: Dulbecco’s modified Eagle medium
DNA: Deoxyribonucleic acid
DS: Docking strand
DTT: Dithiothreitol
ECM: Extracellular matrix
EDTA: Ethylenediaminetetraacetic acid
EGM: Endothelial growth medium
EMT: Epithelial-to-mesenchymal transition
FACS: Fluorescence-activated cell sorting
FBS: Fetal bovine serum
Fc: Fragment crystallizable
FDA: Food and Drug Administration
FDR: False discovery rate
FFPE: Formalin-fixed paraffin-embedded
FFT: Fast Fourier transform
GLMM: Generalized linear mixed model
GO: Gene Ontology
GO-BP: Gene Ontology Biological Process
GSEA: Gene set enrichment analysis
HEPES: 4-(2-hydroxyethyl)-1-piperazineethanesulfonic acid
HRP: Horseradish peroxidase
HUVEC: Human umbilical vein endothelial cell
IACUC: Institutional animal care and use committee
IDT: Integrated DNA Technologies
IF: Immunofluorescence
IS: Imaging strand
LN: Lymph node
LSI: Latent semantic indexing
M-CSF: Macrophage colony-stimulating factor
McDC: Monocyte/classical dendritic cell
MOI: Multiplicity of infection
MSI: Microsatellite instability
MSS: Microsatellite stable
NaOH: Sodium hydroxide
NEAA: Non-essential amino acids
nt: Nucleotide
OCT: Optimal cutting temperature
OHSU: Oregon Health & Science University
PBS: Phosphate-buffered saline
PBMC: Peripheral blood mononuclear cell
PCA: Principal component analysis
PCL: Photocleavable linker
PDAC: Pancreatic ductal adenocarcinoma
PDL: Poly-D-lysine
PDMS: Polydimethylsiloxane
PFA: Paraformaldehyde
PMSF: Phenylmethylsulfonyl fluoride
qRT-PCR: Quantitative reverse transcription polymerase chain reaction
RNA: Ribonucleic acid
RNase: Ribonuclease
RFP: Red fluorescent protein
RIPA: Radioimmunoprecipitation assay
RPM: Revolutions per minute
scATAC-seq: Single-cell assay for transposase-accessible chromatin sequencing
scRNA-seq: Single-cell RNA sequencing
SD: Standard deviation
SEM: Standard error of the mean
shRNA: Short hairpin RNA
SMART-Seq: Switching mechanism at 5′ end of RNA template sequencing
TBST: Tris-buffered saline with Tween 20
TIFF: Tagged image file format
Tn5: Transposase 5
Tris-HCl: Tris(hydroxymethyl)aminomethane hydrochloride
t-SNE: T-distributed stochastic neighbor embedding
UMAP: Uniform manifold approximation and projection
UV: Ultraviolet

## Methods

### Tumor-macrophage hybrid cell identification and analysis in publicly available scRNA-sequencing

#### Single-Cell Data Preprocessing

scRNA-seq data of primary and metastatic CRC patient samples were obtained from a publicly available dataset.^55^ Count matrices were imported using Read10X_h5() and converted into Seurat v5 objects with default thresholds (min. 3 cells per gene, min. 200 features per cell).^90,91^ Metadata files containing cell-level annotations (e.g., sample origin, patient ID, and original cell type) were matched to barcode identifiers and integrated using AddMetaData(). The epithelial and non-epithelial objects were merged using merge() to generate a combined dataset. Quality control was assessed by visualizing metrics such as gene count (nFeature_RNA), transcript count (nCount_RNA), and mitochondrial gene percentage (percent.mt) using VlnPlot(). The merged object was then normalized (NormalizeData()), variable features were identified (FindVariableFeatures()), data were scaled (ScaleData()), and dimensionality reduction was performed via principal component analysis (PCA). Nearest-neighbor graphs and clusters were generated using FindNeighbors() and FindClusters() (resolution = 2.0), and a UMAP embedding was computed with RunUMAP() (dims = 1:30).

#### Hybrid Cell Classification

Hybrid tumor-immune cells were defined based on co-expression of epithelial and immune markers within the RNA assay. Epithelial identity was assigned to cells expressing *EpCAM*, *CDH1* (E-cadherin), and pan-cytokeratin (including *KRT2–5, KRT7–8, KRT14–16, KRT19*), consistent with AE1/AE3 immunoreactivity. Immune identity was assigned based on expression of canonical markers including *PTPRC* (*CD45*), and macrophage markers *CD14, CD68, CD163*, and *ITGAM*. Cells expressing at least one epithelial marker and one macrophage marker were classified as hybrids using WhichCells(). Subsets of hybrids enriched for macrophage lineage markers were annotated as “tumor-macrophage hybrids.” Hybrid classifications were stored in the metadata under hybridcells and counts per cell type and patient were tabulated using table().

#### Doublet Detection and Filtering

To evaluate doublet scores for hybrid cell populations and remove potential doublets, cells were scored using the scds package with three methods: cxds(), bcds(), and the cxds_bcds_hybrid() model based on prior work.^58,92,93^ Hybrid scores were assigned to metadata (cxds-bcds-score, cxdsscore, bcdsscore) and visualized using violin plots (VlnPlot()). Cells with cxds-bcds-score ≥ 0.5 were removed from downstream analyses. Additional doublet scores were calculated using scDblFinder (Supplemental Figure 1) based on our prior work in uveal melanoma.^54^ Seurat object filtering and visualization steps ensured consistency across doublet detection methods. The updated Seurat object excluding high-scoring doublets was used for all downstream hybrid analyses.

#### Differential Expression and Pathway Analysis

Differentially expressed genes (DEGs) were identified using Seurat’s FindMarkers() function with a minimum detection threshold of 25% (min.pct = 0.25). Comparisons were made between hybrid cells and other cell types (epithelial, macrophage-like), as well as between hybrid cells from tumor and lymph node (LN) sites. Top up- and down-regulated genes were visualized using heatmaps (DoHeatmap()) and dot plots (DotPlot()), with log₂ fold changes used to rank genes. For pathway analysis, gene set enrichment analysis (GSEA) was performed using the clusterProfiler package^94^ (gseGO()) with the Gene Ontology Biological Process (GO-BP) database. Gene symbols were converted to Ensembl IDs via org.Hs.eg.db::mapIds(), and redundant gene entries were collapsed by retaining those with the highest absolute log fold change. GSEA plots and enrichment maps were generated using dotplot() and emapplot() from the enrichplot package.

#### Hybrid Cell Subsetting and Clinical Correlation

Clinical metadata including sex, age, tumor stage, MSI/MSS status, and mutational status (e.g., KRAS, BRAF, TP53, APC) were integrated into the object and stored in new metadata columns. Hybrid cell counts were assessed for all available clinical metadata and provided in Supplemental Figure 1. Hybrid cell analyses were restricted to matched tumor and lymph node patient samples (n = 7) where applicable using subset() by patient.ID in Seurat. RUNX1 pathway activity was examined by comparing expression of key regulators (*RUNX1, SPI1, THBS1, LGALS3, PF4, GRN*) in hybrid vs. epithelial or macrophage populations and across cancer stage. Group comparisons were performed using two-sample t-tests and plotted using VlnPlot().

### Generation of in-vitro derived fusion hybrids and single cell clones

MC38-H2B-RFP colorectal cancer cells were co-cultured with bone marrow-derived macrophages isolated from β-actin-GFP C57BL/6 mice to generate spontaneous hybrid cells as previously described.^86^ Briefly, bone marrow cells were harvested from femurs and tibias of a β-actin-GFP C57BL/6 female mouse and were cultured in high glucose DMEM supplemented with 15% FBS, 1% Penicillin/Streptomycin, and 25 ng/mL recombinant M-CSF to promote macrophage differentiation over 4 days. After differentiation, cells were plated at a 2:1 ratio with MC38 cells and co-cultured for 4 days in high glucose DMEM supplemented with 10% FBS, 1% NEAA, 1% HEPES, 1% Penicillin/Streptomycin, and 5 ng/mL recombinant M-CSF. Double-positive GFP+/RFP+ cells were isolated via FACS using a BD influx and processed either immediately for scRNA and scATAC sequencing as described in more detail below (freshly fused/nascent; hybrid-M) or expanded through serial passaging and purity sorting for GFP+/RFP+ cells (hybrid-T). To generate single cell clones, sorted single hybrid cells were deposited into 96-well plates (H1-H12) and wells with single cells were annotated 30 minutes after FACS sorting and cell growth was assessed twice weekly until single cell colonies reached 70% confluence and were further expanded and frozen down for future studies. Tumor cell controls (RFP+) and macrophages (GFP+) were also sorted at the same time using the same methods and processed for scRNA and scATAC studies described below. After sorting, hybrid cells and MC38 tumor cells were grown in high glucose DMEM media supplemented with 10% FBS, 1% NEAA, 1% HEPES, and 1% Penicillin/Streptomycin. This cell culture growth media was used for all experiments unless otherwise noted in the individual experimental methods below. All experiments were completed for cell lines between passages 8-15.

### Phenotypic evaluation of hybrid single cell clones

#### qRT-PCR

Total RNA was extracted using the RNeasy Plus Mini Kit (Qiagen). RNA integrity was assessed with a Nanodrop. cDNA was synthesized using the High-capacity cDNA synthesis kit (Applied Biosystems). qRT-PCR was performed with SYBR Green PCR Master Mix and primer sets from Table 4 on a CFX Opus 384 (Bio-Rad). Relative gene expression was calculated using the ΔΔCt method, using GAPDH as the internal reference gene. Results are mean ± SEM from triplicate analyses averaged from three independent biological replicates. Heatmaps and hierarchical clustering were generated using the pheatmap package in R.

**Table 4:**
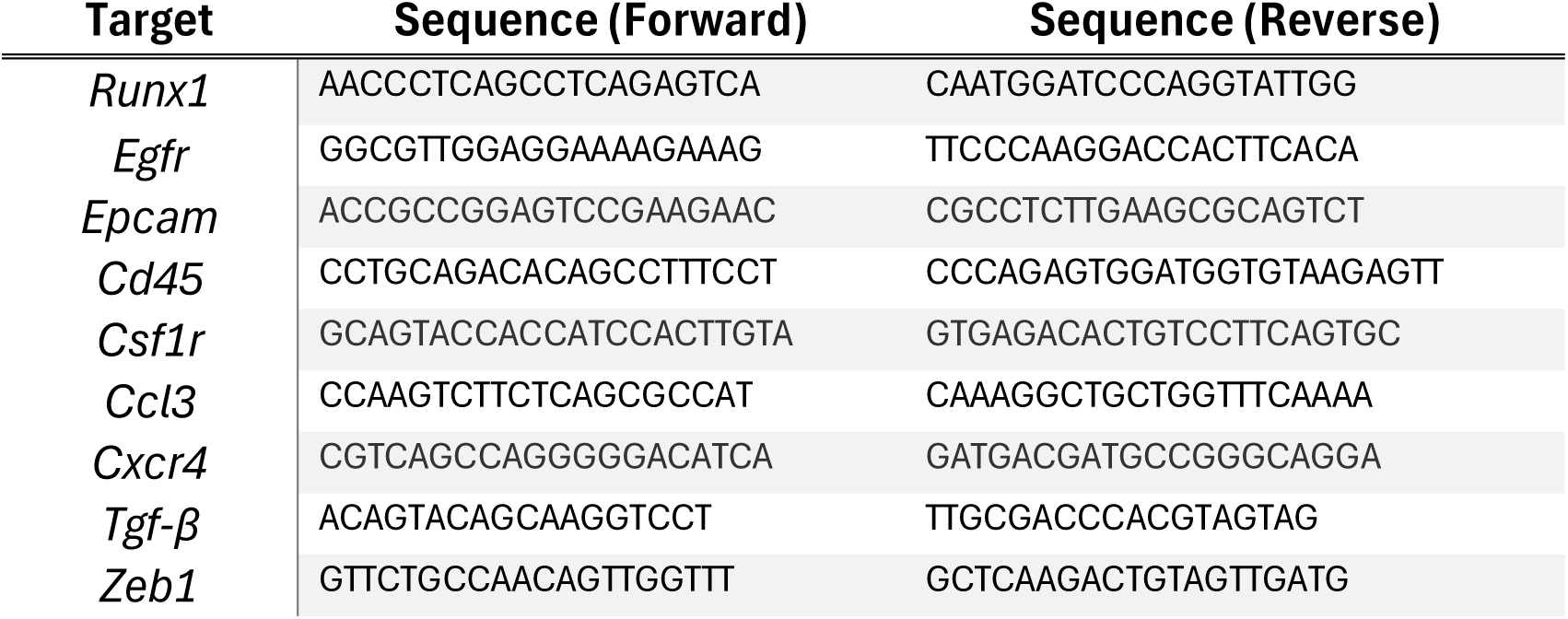
Primer sets for qRT-PCR.

#### Proliferation

To assess cell proliferation, cells were plated in 96-well plates at equal density and monitored over a 72-hour period using the IncuCyte® Live-Cell Analysis System (Sartorius). Phase contrast images were acquired every hour in each well using a 10× objective. Proliferation rates were quantified using the percentage of phase object confluence over time to calculate doubling time (graphed as “1/doubling time”) using IncuCyte’s integrated image analysis software. All experiments were conducted in triplicate, with biological replicates, and data were normalized to initial confluence at time zero.

#### Chemotaxis

Cells were seeded at equal density in the upper chambers of IncuCyte® ClearView 96-well Chemotaxis Plates (Sartorius) in serum-free growth media. The lower wells contained growth media supplemented with 10% fetal bovine serum (FBS) as a chemoattractant. The plate was imaged every hour for 60 hours using the IncuCyte® Live-Cell Analysis System with a 10× objective, capturing phase contrast images of cells migrating through the membrane to the underside of the upper chamber. Cell migration was quantified using IncuCyte’s integrated chemotaxis analysis module, measuring cell count in the lower focal plane over time, normalized to the initial top value count. Each condition was run in at least triplicate wells per experiment, with 2 or more biological replicates for both the hybrid single cell phenotyping studies (Figure 2) and Runx1 shRNA studies (Figure 5).

#### ECM collagen invasion on-a-chip

To establish the invasion on-a-chip, we used an identTx3 microfluidic device (Mattek). The device contains a central channel (1.30 mm in width and 0.25 mm in height) separated by pillars from two adjacent parallel channels (0.5 mm in width and 0.25 mm in height). To improve collagen adherence, the central chamber of the devices was coated with 1 mg/mL of poly-D-lysine (PDL) (Gibco) for 3 hours at 37 °C, then washed with ultrapure water and allowed to dry overnight inside the biosafety cabinet. Next, we prepared collagen hydrogel by mixing 833 μL of acid-solubilized type I collagen from rat tail (3 mg/mL, Gibco) with 100 μL of 10x PBS, 20 μL of 1 M NaOH, and 47 μL of growth media without serum to achieve a working pH of 7.2 to 7.4. Subsequently, 10 μL of collagen was pipetted into the central channel carefully to avoid bubbles. Devices were maintained in the incubator at 37°C for 45 minutes to allow collagen fibrillogenesis, then the lateral channels were filled with media to prevent collagen dehydration. On the next day, cells (MC38 tumor, H11, H4, H3 or H7) in serum-free growth media were seeded in one of the lateral channels of the device while on the other channel, growth media with 10% FBS was placed to act as a chemoattractant for the cells. The devices were then placed in the incubator to allow cells migration into the collagen gel. Three images spanning across each channel were acquired every 24 hours for three days using a Nikon spinning disk live-cell imaging system with z-stack to assess cell migration in x,y and z planes using a 10x objective. After 72 hours, the cells were fixed and stained with DAPI and imaged using the same microscope and imaging parameters. The number of invading cells were quantified by manual annotation in a blinded observer using Zen Blue software (Carl Zeiss Microscopy). Data presented are the total cell counts from 2 biological replicates and 2 technical replicates, n=4 per cell line.

#### Vasculature invasion on-a-chip

##### Fabrication of the microfluidic device

A CAD program (Autodesk Fusion 360 v.2606.1.36) was used to design a mold composed of two reservoirs connected by a central channel and a chamber as published.^95^ We then used a three-dimensional (3D) printer (CADworks3D μMicrofluidic printer, Profluidics 285D) and resin (Master Mold resin, CADworks) to print the positive molds. Printed micromolds were cleaned in methanol in three rinses of 2 minutes each under agitation, subsequently were cast with PDMS (Polydimethylsiloxane - Sylgard 184, Dow-Corning), and left to cure overnight at 80°C, as previously described.^95,96^ Next, PDMS was removed from the resin mold, and two reservoirs were prepared using a 5-mm biopsy punch for media, 1-mm biopsy punches for the collagen loading ports and 2.5 mm punch on top of the central chamber where collagen will be placed. Molds were cleaned with ethanol and immediately plasma bonded to a glass coverslip. Assembled devices were autoclaved, then treated with 1% (w/v) glutaraldehyde (Sigma-Aldrich) for 15 minutes, rinsed three times with distilled water (DIW), and left overnight in DIW to remove any trace of glutaraldehyde. To mold cylindrical channels, sterile 160 μm diameter acupuncture needles were immersed in a 0.1% bovine serum albumin (BSA) solution for at least 30 minutes and then inserted into the central channel of the devices ∼200 μm above the glass coverslip surface. Rat tail collagen type I (3.0 mg/mL, Gibco) was prepared according to published protocol.^95^ Briefly, a stock solution of collagen type I (collagen I, rat tail Gibco, 3 mg/mL) was prepared on ice by mixing 833 μL of collagen with 100 μL of 10x PBS, 20 μL of 1 M NaOH, and 47 μL of EGM to reach a working pH of 7.2-7.4. The final solution was immediately pipetted into the middle chamber of the device and allowed to polymerize at 37°C.^95^ To prevent collagen dehydration, after 1 hour, the main reservoirs and the top of the collagen hydrogel were filled with EGM, then devices were returned for incubation overnight. On the following day, the needles were carefully removed with a pair of tweezers, and the cell medium (EGM) was replaced by fresh medium. Subsequently, the devices had their reservoirs filled with fresh EGM and were placed in the incubator at 37°C and 5% CO2 overnight before cell seeding.

##### Cell culture

Human umbilical vein endothelial cells (HUVECs expressing green fluorescent protein (GFP-HUVECs, Lonza, Basel, Switzerland) were cultured in a supplemented media (EGM-2 with bullet kit, Lonza). Human bone marrow mesenchymal stem cells (hMSCs, RoosterBio, MD) were cultured in α–minimum essential medium (Gibco) with 10% FBS and 1% penicillin/streptomycin. Cell media was changed every other day, and cells were passaged when reaching a confluency of 80-90%. HUVECs at passages 4-6 and hMSCs at passages 2-4 were used for all the experiments. All cells were maintained in a humidified incubator (5% CO_2_ and 37°C).

##### Cell seeding

For seeding, GFP-HUVECs and hMSCs were detached using TrypLE (Gibco, FisherScientific), counted, and mixed at a 4:1 ratio according to previous publications^95^ in a cell density of 6×10^6^ cells/mL. Subsequently, the cell medium was removed from the reservoirs, and 25 μL of the cell suspension was added into one reservoir. The devices were flipped upside down, placed in the incubator for 5 minutes, seeded again, and left upside down in the incubator for another 5 minutes. Until the entire extension of the collagen channel had cells attached, we repeated the seeding, flipping the chip as needed. Next, the devices were placed in the incubator for 30 minutes under static conditions. Afterward, the devices were transferred to the two-dimensional (2D) rocker (BenchRocker) inside the incubator as published.^95,96^ After 24 hours, vasculature was formed and 4 x10^5^ hybrid cells or unfused parent cells were seeded on the central reservoir of the devices, on top of the collagen so that cells were 800 µm away from the vascular channel. The devices were maintained in a humidified incubator (5% CO_2_ and 37°C) for 24 hours and fixed with 4% PFA (v/v) in PBS for 30 minutes.

##### Cell staining and imaging

Cells were permeabilized with 0.1% (w/v) Triton X-100 for 10 minutes and blocked with 1.5% (w/v) BSA for 1 hour under agitation. After washing with PBS, samples were incubated with one of the following primary antibodies (anti-PECAM-1, rabbit anti-human, cat. no. AB32457, Abcam, 1:100; or anti-NG-2, mouse anti-human, cat. no. 14-6504-82, Invitrogen, 1:200) overnight at 4°C. Samples were washed with PBS and incubated with secondary antibody (goat anti-mouse Alexa Fluor 555, Invitrogen, 1:250; goat anti-rabbit Alexa Fluor 647, Invitrogen, 1:250) overnight at 4°C under agitation. This was followed by rinsing in 0.1% PBS, staining of the nuclei using NucBlue [Fixed Cell ReadyProbes, 4′,6-diamidino-2-phenylindole (DAPI), ThermoFisher], and staining of actin with ActinGreen 488 (ReadyProbes, ThermoFisher) for 1 hour at 37°C under agitation.

Samples were imaged using a confocal microscope (Zeiss, LSM 880, Germany) with a 10× objective (numerical aperture, 0.45; Zeiss, Plan Apochromat). The depth of imaging was 100 to 400 μm, split in at least 100 *z*-stacks. *z*-stacks were converted into TIFF files or 3D images using Zen Blue software (Carl Zeiss Microscopy) or Imaris software (version 9.1, Bitplane, Oxford Instruments, Zurich, Switzerland). The number of migrating cells were counted using Imaris.

#### Protease activity

##### Protease Probe Synthesis and Labeling

Protease target probes (ACE2, CATB, MMP14, MTP, PLASM, USP15) were synthesized at >98% purity with N-terminal acetylation and C-terminal amidation (GenScript). Probes were conjugated to the fluorescent reporter BDP FL NHS ester (Lumiprobe, Cat. #41420) and aliquoted for single-use storage. Immediately prior to each assay, aliquots were thawed on ice and protected from light.

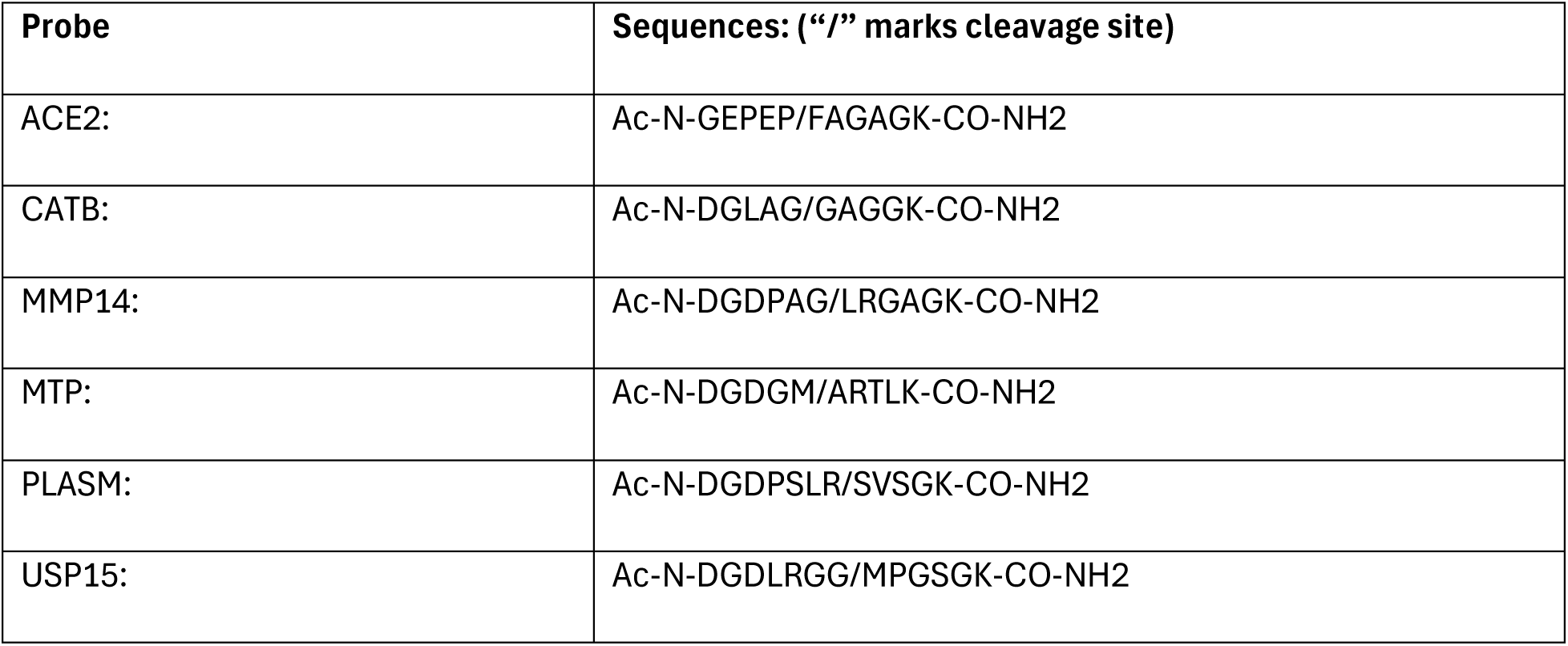

| **Probe** | **Sequences: (“/” marks cleavage site)** |
| --- | --- |
| ACE2: | Ac-N-GEPEP/FAGAGK-CO-NH<sub>2</sub> |
| CATB: | Ac-N-DGLAG/GAGGK-CO-NH<sub>2</sub> |
| MMP14: | Ac-N-DGDPAG/LRGAGK-CO-NH<sub>2</sub> |
| MTP: | Ac-N-DGDGM/ARTLK-CO-NH<sub>2</sub> |
| PLASM: | Ac-N-DGDPSLR/SVSGK-CO-NH<sub>2</sub> |
| USP15: | Ac-N-DGDLRGG/MPGSGK-CO-NH<sub>2</sub> |

##### Protease Reaction Setup

Reactions were performed in 96-well V-bottom plates. Each well contained 2 µL of 50 mM CaCl₂, 4 µL of peptide working stock, and 4 µL of lysate, for a final reaction volume of 10 µL per well. The final probe concentration was 200 µg/mL with 10 mM CaCl₂. Plates were sealed, wrapped in foil, and incubated on an orbital shaker at room temperature for 60 minutes.

##### Sample Preparation and Gel Electrophoresis

Following incubation, 2 µL of 60% sucrose was added to each well. Samples (10 µL) were loaded into 20% acrylamide TBE gels, each in triplicate, and electrophoresed at 300 V for 60 minutes under inverted polarity. Gels were rinsed with deionized water prior to imaging.

##### Imaging and Quantification

Fluorescent signal was captured on an iBright 1000FL imaging system (ThermoFisher Scientific) using the following settings: 1X digital zoom, 1.5X optical zoom, and focus level 370. Images were acquired at exposure times of 20, 50, and 100 ms. Band intensities were quantified using iBright Analysis Software (ThermoFisher Scientific).

#### *In vivo* tumor models

All animal procedures were conducted in accordance with protocols approved by the Oregon Health & Science University (OHSU) Institutional Animal Care and Use Committee (IACUC). Mice were maintained in a specific pathogen-free facility under a 12-hour light/dark cycle with unrestricted access to standard rodent chow (5001, PMI Nutrition International, Richmond, IN) and water. Experimental cohorts included adult mice aged 8 to 12 weeks, with balanced representation of both sexes, and littermate controls were used where applicable. MC38, H11, and H7 cells were injected into C57BL/6 mice subcutaneously at a dose of 50,000 cells diluted 1:1 in sterile PBS/Matrigel. Tumor volume using calipers and body weight were measured twice weekly until tumor endpoint (reaching a maximum tumor size of 1200mm^3^). At that time, mice were euthanized, and tumors were harvested for FFPE and OCT embedding.

##### RUNX1 Inhibition

For RUNX1 inhibition studies, mice bearing H11 tumors were randomized to treatment once tumors reached approximately 300 mm³. Mice were treated with the RUNX1 inhibitor Ro5 -3335 (5 mg/kg) or vehicle control via intraperitoneal injection every 3 days for a total of three doses. Tumor growth was monitored as described above, and circulating hybrid cells were assessed at endpoint via immunofluorescent staining of isolated PBMCs adhered to slides as detailed below.

##### Sample Preparation, Staining, Imaging and Quantification

PBMCs were isolated via Ficoll density centrifugation, adhered to poly-D-lysine–coated glass slides, and fixed in 4% paraformaldehyde (PFA) prior to staining as previously described.^97^ PBMC cell slides and FFPE tumor sections were washed 3×5 minutes with PBS and incubated in blocking buffer (2.5M CaCl_2_, 1% TritionX-100, 1% BSA in PBS) for 30 minutes at room temperature. Primary antibodies targeting GFP (GFP-1020, Aves Labs) and RFP (5f8, ChromoTek) were applied at a 1:500 dilution in blocking buffer overnight at 4°C. The next day, secondary fluorescent antibodies Donkey Anti-Chicken Alexa Fluor 488 (Jackson ImmunoResearch, 703-546-155) and Donkey Anti-Rat Cy3 (Jackson ImmunoResearch, 712-165-153) were applied at a 1:500 dilution, incubated for 45 minutes at room temperature, and counterstained with DAPI. Whole slides were imaged using an AxioScan.Z1 digital slide scanner (Zeiss) equipped with a Colibri 7 light engine. Exposure times were adjusted per fluorophore and antibody to optimize signal detection and avoid saturation and were determined using negative staining controls for each round. Imaging was performed using a 20x Plan-Apochromat 0.8 NA objective and stitched using Zen Blue software (Carl Zeiss Microscopy). GFP^+^ cells were quantified in QuPath^98^ (Version 0.6.0-rc5). Cells were segmented using the InstanSeg extension, and thresholds for GFP positivity were determined from a paired secondary antibody-only control section for each animal. All GFP^+^ cells were manually reviewed to exclude artifact signal. Data are reported as GFP^+^ cells per 50,000 nuclei. For the FFPE tumor sections, fluorescence reporter signal was quantified using FIJI (Version 2.16/1.54p). Acquisition settings were held constant across all samples. Thresholds for GFP and RFP channels were established using A431 tumors lacking both GFP and RFP reporter expression and applied uniformly to all samples. Reporter signal was quantified as area fraction above threshold within manually defined tissue masks.

In vivo tumor growth experiments were performed in biological triplicate. Cohort sizes varied due to differences in tumor engraftment rates across cell lines. Data shown represent the largest experimental cohort with a balanced male-to-female ratio and are representative of findings observed across independent experiments. RUNX1 inhibitor studies were performed in biological duplicate. RUNX1 inhibitor tumor growth studies were performed in biological duplicate with n=10 mice per treatment group with a balanced male-to-female ratio (5M/5F). Analysis of CHCs after treatment with RUNX1 inhibitor was limited by compromised peripheral blood samples (n=5 per treatment group).

### Runx1 shRNA studies

#### Cell culture

MC38-RFP tumor cells and two MC38xY01-derived hybrid clones (Clone 7 and Clone 11) were cultured in high glucose DMEM medium supplemented with 10% FBS, 1% NEAA, 1% HEPES and 1% penicillin-streptomycin. Neomycin (G418, Sigma Aldrich) titration was performed prior to transduction to determine the minimum concentration required to eliminate non-transduced cells. Cells were seeded in 24-well plates and treated with a dilution series of neomycin (G418). At 1000 µg/mL, all cells were killed within 4 days. MC38 tumor cells showed slightly higher sensitivity and were also maintained under the same neomycin concentration for consistency.

#### Lentiviral transduction

Lentiviral shRNA particles targeting RUNX1 (TRCN0000084810 and TRCN0000084812) and a non-targeting control (TRCN0000072229) (Sigma Aldrich) were used at calculated multiplicities of infection (MOIs) based on viral titers and cell counts (Table 5). Cells were seeded into 24-well plates at densities to reach target densities for MOI calculation the following day. Immediately prior to transduction, media was replaced with fresh growth medium containing 8 µg/mL polybrene (hexadimethrine bromide), except for one control well that received polybrene-free medium to monitor potential toxicity. The appropriate amount of viral supernatant was added directly to each well based on calculated MOI of 20 viral particles per cell and placed back into the incubator overnight. The next day, viral-containing media were aspirated and replaced with fresh growth medium for 24 hours before starting selection with neomycin (G418) containing media. Transduced cells were selected by growing in media containing 1000 µg/mL neomycin. Brightfield images were captured at each media change to document cell viability and morphology. Due to insufficient knockdown efficiency observed in preliminary experiments, a second transduction was performed using the same procedure. This repeat transduction was carried out after the first round of selection and expansion. Cells were re-seeded at appropriate densities, and transduction was repeated as described above. Selection with 1000 µg/mL neomycin was re-applied and cells were maintained under selection for 72 hours before downstream analyses including qRT-PCR, western blot, chemotaxis, proliferation, ECM invasion, and protease activity as previously described.

**Table 5:**
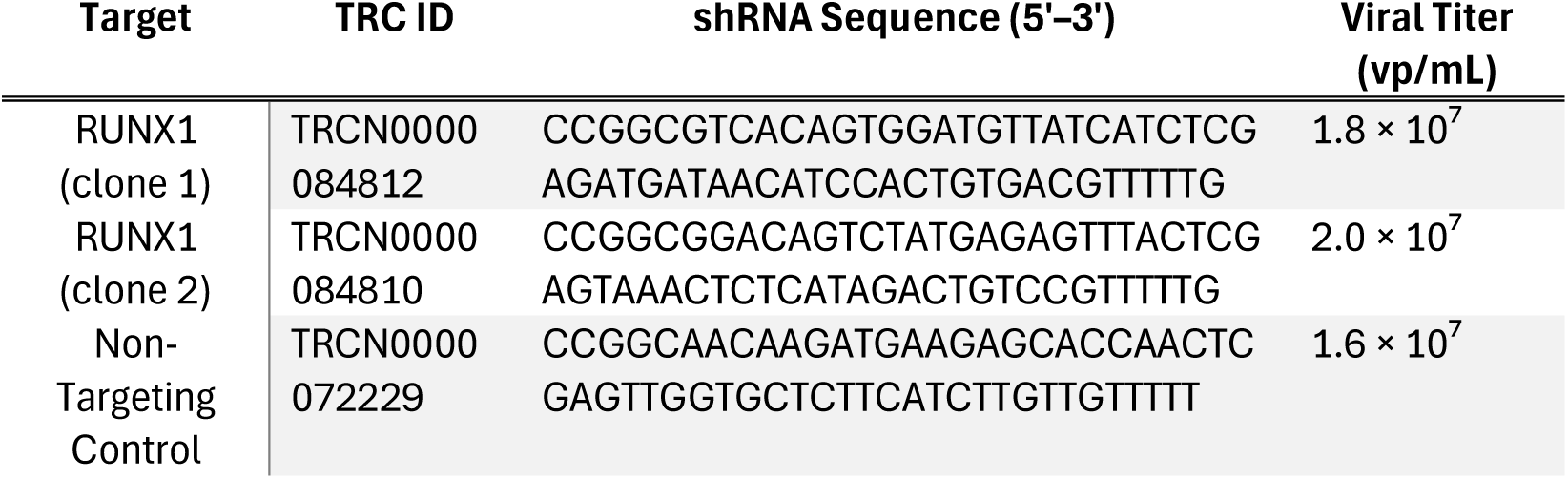
Runx1 lentiviral constructs.

#### Western Blots

Cells were harvested for western blot analysis using trypLE (Gibco, FisherSci) to detach adherent cells. After pelleting cells were washed once with 1X PBS, pelleted and were then either flash-frozen in liquid nitrogen or immediately lysed. Cell lysis was performed using RIPA buffer (ThermoFisher Scientific, 89900) supplemented with 1 mM PMSF and one protease inhibitor mini-tablet (per 7 mL buffer) and phosphatase inhibitor cocktail 2 (Sigma-Aldrich). Lysis buffer was added at 60 μL per 1×10⁶ cells. Samples were incubated on ice for 20 minutes to allow solubilization. Samples were sheared to reduce viscosity via vigorous vortex. Lysates were centrifuged at 15,000 × g for 10 minutes at 4°C, and supernatants were transferred to new tubes. Protein concentration was normalized using a BCA assay. Lysates were mixed with 3× GS loading buffer (6% SDS, 150 mM Tris pH 8.0, 30% glycerol, 0.1% bromophenol blue) containing 1× final concentration of β-mercaptoethanol (BME) and heated at 95°C for 5 minutes. Samples were loaded into Criterion TGX gels (Bio-Rad) and electrophoresed at 130–150 V for ∼1 hour in 1× running buffer. Proteins were transferred to PVDF membranes (activated in ethanol) using the Bio-Rad Trans-Blot Turbo system according to manufacturer-specified settings based on protein molecular weight. Membranes were blocked for 5 minutes at room temperature in EveryBlot blocking buffer (Bio-Rad, 12010020). Blots were incubated overnight with primary antibodies at 4°C (RUNX1 (19555-1-AP, ProteinTech) 1:1000; ACTIN (2066, Sigma) 1:100) in TBST, followed by 3 x 5-minute TBST washes. Secondary antibody (goat anti rabbit HRP 1:2000) was applied for 1–2 hours at room temperature, followed by additional TBST washes (1-2 hours). Signal detection was performed using SuperSignal West Pico PLUS or Femto chemiluminescent substrates (ThermoFisher Scientific), and membranes were imaged using an iBright FL digital imaging system (ThermoFisher Scientific). Bands were quantified using iBright analysis software (ThermoFisher Scientific).

#### Immunofluorescent staining and imaging

Cells were grown on ibidi chamber slides and fixed using 4% PFA for 15 minutes at room temperature once desired confluency was reached (∼70% confluent). After fixation, cells were washed 3×5 minutes with PBS and incubated in blocking buffer (2.5M CaCl_2_, 1% TritionX-100, 1% BSA in PBS) for 30 minutes at room temperature. Primary antibody targeting RUNX1 (RUNX1 PA5-19638, 1:800) was applied diluted in blocking buffer overnight at 4°C. The next day, secondary fluorescent antibody was applied at 1:500 dilution and incubated for 1 hour at room temperature and counterstained with DAPI. Cells were imaged using an inverted epifluorescence microscope (Axio Observer, Zeiss) at various magnifications (10x, NA 0.45; 20x, NA 0.8, 40x NA 1.3). All images shown are representative of three or more images collected per cell line, per condition, in biological triplicate. Quantification of fluorescent intensity was analyzed using Zen Blue software (Carl Zeiss Microscopy).

### cyCIF CRC patient matched tumor and peripheral blood

#### Antibody Generation

Antibodies were prepared using barcoding technologies as previously described.^99–102^ In brief, each antibody was site-specifically conjugated to a 28-nucleotide (nt) single-stranded DNA docking strand (DS) using the SiteClick™ Antibody Azido Modification Kit (ThermoFisher Scientific), which targets the Fc region. For marker detection, 26 nt complementary imaging strands (IS) labeled with fluorophores and photocleavable linkers (PCLs) at both the 5′ and 3′ termini were hybridized to the docking strands. All oligonucleotides were synthesized by Integrated DNA Technologies (IDT, Coralville, IA). A complete list of antibodies is provided in Table 3.

#### Staining Protocol & Signal Removal

cyCIF was performed on formalin-fixed paraffin-embedded (FFPE) tumor tissue sections and corresponding peripheral blood mononuclear cells (PBMCs) from patient-matched specimens, as previously described and detailed in Table 2.^45,99–102^ FFPE tumor sections (5 µm) were first deparaffinized with xylene and rehydrated through a graded ethanol series. Antigen retrieval was conducted by incubation in citrate buffer (pH 6.0) for 30 minutes at 100°C, followed by rinsing in heated deionized water and a 10-minute incubation in Tris-HCl buffer (pH 8.0) at 100°C. Slides were then cooled to room temperature and washed in PBS.

PBMCs were isolated via Ficoll density centrifugation, adhered to poly-D-lysine–coated glass slides, and fixed in 4% paraformaldehyde (PFA) prior to cyCIF staining. Cells were rehydrated in 3×5 minute PBS washes. Tumor and PBMC slides were blocked for 30 minutes at room temperature in PBS containing 2% bovine serum albumin (BSA), 0.5% dextran sulfate, and 0.5 mg/mL sheared salmon sperm DNA. Ab-oligo conjugates targeting specific biomarkers (Table 3) were diluted in blocking buffer and applied to the tissue and blood samples in a single-step staining protocol (separated into two rounds of primary antibody incubation (after R0 and after R5) as the volume of 34 antibodies left very little blocking buffer added to dilute to intended concentrations). Following overnight incubation at 4°C, unbound Ab-oligos were removed by washing 3×5 minutes using 2X saline-sodium citrate buffer (SSC), and samples were post-fixed with 2% PFA for 10 minutes. Samples were then incubated with imaging strands (IS) specific to the DNA docking sequences on the antibodies for 45 minutes, washed 3×5 minutes in 2x SSC and counterstained with DAPI for 10 minutes. Coverslips were mounted using fluoromount-G (ThermoFisher Scientific). Three to four ISs were applied per imaging round, each labeled with a distinct fluorophore, DAPI (Zeiss 96 HE), Alexa Fluor 488 (Zeiss 38 HE), AF555 (Zeiss 43 HE), AF647 (Zeiss 50), and AF750 (Chroma 49007 ET Cy7).

#### Image Acquisition

Whole slides were imaged using an AxioScan.Z1 digital slide scanner (Zeiss) equipped with a Colibri 7 light engine. Exposure times were adjusted per fluorophore and antibody to optimize signal detection and avoid saturation and were determined using negative staining controls for each round. Imaging was performed using a 20x Plan-Apochromat 0.8 NA objective and stitched using Zen Blue software (Carl Zeiss Microscopy).

#### Fluorescent Signal Removal

After each round of imaging, fluorescence signals were removed by UV treatment for 15 minutes to enable additional rounds of staining. The cycle of IS hybridization, imaging, and UV-based fluorophore cleavage was repeated until all desired markers were imaged. Round 0 (R0) images were acquired following DAPI staining and prior to antibody application to assess background autofluorescence using the 488 channel. Two tumor sections from different tumor regions and one peripheral blood specimen (∼1.5 mL) were analyzed per patient.

#### Image analysis

For Ab-oligo validation, Zen Blue software (Carl Zeiss Microscopy) was used for image visualization.

For cyclic immunofluorescence, images from cyclic staining rounds were registered using ASHLAR feature-based image registration.^103^ Cells were segmented using QiTissue default segmentation parameters for PBMC slides and using MESMER segmentation for tumor tissues.^104^ The edges of the tissue sections and wells were excluded from analysis as well as areas with imaging artifacts (e.g., out of focus, bubbles). Using QiTissue, cells with high autofluorescence were removed from analysis based on the whole cell average fluorescence from the round 0 background channel. Hybrids were defined as co-expressing CD45 and at least one of the epithelial markers: panCK, ECAD, or EpCAM. Positive expression was defined in QiTissue as a whole cell intensity average value above the positive staining threshold that was determined by normalizing to an unstained control for each patient. Average cell intensity values were extracted for each antigen in the phenotyping panel for each identified hybrid cell and circulating tumor cell.

Extracted average cell intensity features for all cells, including CHCs and CTCs, were compiled into a unified feature table and normalized per marker using UniFORM feature-level normalization method.^105^ UniFORM aligns marker intensity distributions across samples via FFT-based cross-correlation while preserving distribution shapes and positive population proportions – effectively addressing the right-skewed, heterogeneous nature of multiplexed imaging data. Following normalization, CHCs and CTCs were isolated and merged into a combined dataset and then the normalized CHC/CTC dataset was subjected to Leiden clustering to identify distinct subpopulations. Differential biomarker expression was assessed across clusters and visualized using heatmaps. Expression levels of key markers (e.g., RUNX1, SPI1, THBS1) were visualized, stratified by tumor stage and cell type. Expression differences by stage were evaluated using two-tailed t-tests comparing expression levels between stages (1 vs. 2, 2 vs. 3, and 1 vs. 3).

### Single cell omics in-vitro derived hybrid cells & CRC patient hybrid scRNA sequencing

#### Flow cytometry

All FACS used in these studies was performed on a BD Influx.

For *in vitro* derived hybrid cells GFP and RFP fluorescence was used to isolate double positive hybrid cells. Cells were also stained for viability using DAPI. Tumor cell controls (RFP+) and macrophages (GFP+) were also sorted at the same time using the same methods and processed for scRNA and scATAC studies described below.

For patient samples, PBMCs were isolated using Ficoll-Paque density gradient centrifugation as previously described.^20,45^ Briefly, peripheral blood samples were collected from patients using heparinized vacutainer tubes (BD Biosciences, Franklin Lakes, NJ, USA) and diluted at a 1:2 ratio with phosphate-buffered saline (PBS; 1.37 M NaCl, 27 mM KCl, 0.1 M Na₂HPO₄, 18 mM KH₂PO₄, pH 7.4). 12 mL of Ficoll was underlaid beneath the diluted blood and centrifuged at 800 × g for 20 minutes at room temperature with no brake. The PBMC layer was collected, washed, and resuspended in PBS. Cells were then seeded onto poly-D-lysine–coated slides (1 mg/mL; Millipore, Burlington, MA, USA; Fisher Scientific, Waltham, MA, USA) and incubated at 37°C for 15 minutes to promote adherence. Adherent cells were fixed with 4% paraformaldehyde (PFA) for 5 minutes, permeabilized with 0.5% Triton X-100 (Fisher Scientific, BP151-100) for 10 minutes, and post-fixed with 4% PFA for an additional 10 minutes.

Patient tumor tissue obtained immediately after surgical removal was digested into single cell suspension by finely mincing, followed by enzymatic digestion with Liberase-TH (5 mg/mL in sterile ultrapure H2O) for 30-60 minutes on a stir plate at 300 RPM and 37°C, then filtered using 100 µM cell strainer, pelleted and washed. PBMCs and tumor cells were stained with antibodies targeting EpCAM, ECAD, CD45, and DAPI as previously described.^21^ Briefly, cells were washed and resuspended in FACS buffer [phosphate-buffered saline (PBS), 1.0 mM EDTA, and 5% fetal bovine serum (FBS)]. Cells were incubated in FACS buffer containing Fc Receptor Binding Inhibitor (5 µL5” per 1×10^6^ cells; eBioscience) for 20 minutes on ice. Cells were then incubated in FACS buffer for 30 minutes on ice with CD45-AF488 (1:100; ThermoFisher Scientific), ECAD-AF647 (1:100), EpCAM-AF555 (1:100). DAPI was used to assess cell viability. Single color controls used for gating included EpCAM/Ecad expressing epithelial cell line, and donor PBMCs stained with only CD45-AF488 (1:100). Cells were defined as tumor hybrid cells or circulating hybrid cells by (EpCAM+ and/or ECAD+ and CD45+). Tumor cells (EpCAM+ and/or ECAD+ and CD45−) and immune cells (EpCAM- and ECAD- and CD45+) were also sorted into 16 wells of the 96 well plate containing cell lysis buffer (0.2% TritonX diluted in sterile, ultrapure H_2_O and RNase inhibitor (Takara Bio). Immediately after single cell sorting, plates were sealed, spun down and rapidly frozen and stored at -80°C until further processing for Smart-seq described below. *In vitro* studies data reflect analyses from n=2 hybrid fusion, sorting and downstream 10X sequencing experiments. For patient samples (Table 2), n=3 patients with CRC (2 tumor samples, 3 PBMC samples) were processed for Smart-Seq plate based single cell sequencing, and if substantial hybrid cells obtained, also pooled with tumor cells or PBMCs for 10X genomics sequencing studies described in detail below.

#### Nuclei Isolation for 10x and s3 Libraries

After flow sorting, the nuclei were spun down (5 minutes, 500xg, 4°C) to remove the media supernatant and resuspended it in 1 mL of NIB-Hepes buffer. The cell suspension were then incubated for 5 minutes on ice; and subsequently spun down (5 minutes, 500xg, 4°C). The pellet was resuspended in 1 mL NIB-Hepes, spun again, and resuspended in 1 mL once more before quantification.

#### 10x Genomics scRNA

For the scRNA libraries, isolated nuclei were diluted in 10x Genomics 20x Nuclei Buffer to the desirable targeted cell recovery. The 10x Genomics Single Cell 3’ Gene Expression Kit was then used according to the published protocol.

#### SmartSeq scRNA

For these patient libraries, the nuclei were frozen down in a 96-well plate in lysis buffer as previously described in FACS section above. The published protocol from Takara’s SMART-Seq mRNA Single Cell LP User Manual was followed starting from Step V, First-Strand cDNA Synthesis. The libraries were indexed using the Unique Dual Index Kits, pooled, cleaned, and quantified as described below.

#### ScaleBio Tagmentation for scATAC

To generate these scATAC libraries the s3-ATAC protocol on the iCell8 was leveraged.^106^ The iCell8 instrument was put through standard instrument start up including tip cleaning, stream checks, wash priming, and prechilled the block. Next, the source plate set up was assigned to (Cells – PCR Mix1 – Index 1 (i7 TruSeq) – PCR Mix2 – Index 2 (i5 Nextera) and read in the barcode of the nanochip.

The 100 µM iCell8 TruSeq Primer source plates were prepared. After removing the seal from nanochip and vacuum sealing it into position, the iCell8 first dispensed the i7 and then i5 primers into the chip. Following each dispense the chip was blotted, sealed, and centrifuged (3 minutes, maximum RFC, 4°C) before the next step. During the centrifugation the iCell8 tips were cleaned twice.

The quantified, isolated nuclei were prepared to get them to the appropriate concentration for tagmentation. A master mix of the following (2.67 µL NIB-H, 3 µL 4.3478x TAPs-TD (1M TAPs, 5M KOAc, 1M MgOAc, 1M D-glucosamine, 6.1% DMF, in H_2_O), 0.1 µL pluronic) was made. This master mix was dispensed into a 96-well plate with a multichannel, and to this 5 µL of 500 µM Scale SBS12/A14 TN5 on the Bravo was stamped in. Next 6,000 nuclei were manually pipetted in 4.23 µL NIB-H to the respective wells. We proceeded to tagment at 55°C for 15 minutes. After tagmentation the samples were allowed to cool on ice, before they were pooled, spun down (5 minutes, 500xg, 4°C, with 180° turn) and resuspended in 1 mL TMG Buffer (1.25 mL 4x TAPS Premix, 1.5 mL 50% glycerol, 2.2 mL H_2_O, 50 µL 10% pluronic F-127) to cushion the nuclei and aid in recovery. These TMG washes were repeated three more times, with the final resuspension being in 120 µL iCell8 Loading Buffer (44 µL 4.3478x TAPs-TD, 50 µL TN5 Dialysis Buffer (10 mM Hepes 7.2, 20 mM NaCl, 0.02% TritonX, 2.5% glycerol, 0.2mM DTT), 0.1% 10% pluronic f-127, 401 µL H_2_O). Nuclei were quantified, with a typical recovery at approximately 50% post-tagmentation, and diluted to get 358 nuclei/µL.

Now the tagmented nuclei were read to load to the iCell8 chip. Approximately 35 µL of nuclei were dispensed into each well of the source plate, with 100 µL of cell loading buffer in the control wells. After using the iCell8 to dispense the cells into the chip, it was spun down (10 minutes, maximum RFC, 4°C) while 3 additional tip cleans were run on the iCell8. The chip was incubated at 53.7°C for 15 minutes. After adding 100 µL of PCR master mix (188.4 µL 5x hi GC Buffer, 18.84 µL dNTPs, 14.58 µL Kapa HiFi Polymerase (non-hot start), 316.5 µL H_2_O) to the source plate wells, 150 nL was dispensed into each well of the iCell8 chip. Again, the chip was spun down (10 minutes, maximum RFC, 4°C) while 3 tip cleans were run concurrently. Following this, 12 cycles of sciATAC-P PCR were run with the samples on the iCell8 chip. The library was collected utilizing the iCell8 Collection Kit (Cat. No. 640212) and clean with a standard double sided-SPRI bead protocol prior to quantification.

#### Artificial Doublet Experiment

To generate and assess artificial doublet cell transcriptomes for comparison with hybrid fusions, the Takara’s SMART-Seq Pro application on the ICell8 single-cell system was used (Takara Cat. No. 640257). The experimental design leveraged the previously established distinct fluorescent markers: macrophages expressed GFP, tumors expressed RFP, and hybrid cells co-expressed both fluorophores. The SMART-Seq Pro manual was followed closely, with several key modifications. The ICell8 system dispenses cells according to a Poisson distribution, which can result in imperfect single-cell loading. To increase throughput, the standard protocol involves two consecutive cell dispenses, each followed by imaging to identify wells containing single cells. These wells are then selected for downstream reagent addition. In our modified approach, two separate 384-well source plates were used. For source plate preparation, *in vitro*-derived hybrid cells, cultured MC38 tumor cells, and first-passage macrophages were isolated via flow cytometry as previously described. Cells were stained with DAPI (10 µg/mL) in 2 mL of media on ice and quantified. Two 384-well source plates were prepared: one containing macrophages and hybrids, and the second containing MC38 cells (loaded into wells A1 to D2). During the first dispense, macrophages were loaded into ¾ of the chip and hybrid cells were loaded into the remaining ¼. After imaging, the "Emptywells" filter file was manually edited in the Experiments Folder, changing all 0s to 1s for wells that received macrophages. Further, all hybrid cell wells were set to 1 to prevent further dispensing into those wells. Next, the second source plate was swapped in containing MC38 tumor cells for the second dispense. These cells were only dispensed into the ¾ of the chip that had previously received macrophages, creating artificial doublets. A second round of imaging followed, and the chip was frozen afterward. To identify cell types in each well, the CELLSELECT’s training feature was used to classify wells based on DAPI, GFP, and RFP signals. While high-confidence calls (confidence score ≥ 0.80) were accepted, wells with lower scores were manually reviewed and annotated using predefined criteria due to variability in fluorescence intensity. All subsequent steps—including first-strand cDNA synthesis, cDNA amplification, tagmentation, indexing, and library purification—were performed according to the Takara SMART-Seq Pro manual. Libraries were quantified and sequenced as described in subsequent sections.

| Scan 1 |  | Scan 2 |  | Annotation |
| --- | --- | --- | --- | --- |
| DAPI 0 | GFP 0 | DAPI 0 | RFP 0 | Removed |
| DAPI 1 | GFP 0 | DAPI 0 | RFP 0 | Macrophage |
| DAPI 0 | GFP 1 | DAPI 0 | RFP 0 | Macrophage |
| DAPI 1 | GFP 1 | DAPI 0 | RFP 0 | Macrophage |
| DAPI 0 | GFP 0 | DAPI 1 | RFP 1 | Tumor |
| DAPI 0 | GFP 0 | DAPI 0 | RFP 1 | Tumor |
| DAPI 0 | GFP 0 | DAPI 1 | RFP 0 | Tumor |
| DAPI 0 | GFP 0 | DAPI 2 | RFP 0 | Tumor Doublet |
| DAPI 0 | GFP 0 | DAPI 2 | RFP 1 | Tumor Doublet |
| DAPI 0 | GFP 0 | DAPI 2 | RFP 2 | Tumor Doublet |
| DAPI 1 | GFP 0 | DAPI 1 | RFP 1 | Mac-Tum Doublet |
| DAPI 1 | GFP 0 | DAPI 0 | RFP 1 | Mac-Tum Doublet |
| DAPI 1 | GFP 0 | DAPI 1 | RFP 0 | Macrophage |
| DAPI 1 | GFP 0 | DAPI 2 | RFP 0 | Mac-Tum Doublet |
| DAPI 1 | GFP 0 | DAPI 2 | RFP 1 | Mac-Tum Doublet |
| DAPI 1 | GFP 0 | DAPI 2 | RFP 2 | Removed |
| DAPI 0 | GFP 1 | DAPI 1 | RFP 1 | Mac-Tum Doublet |
| DAPI 0 | GFP 1 | DAPI 0 | RFP 1 | Removed |
| DAPI 0 | GFP 1 | DAPI 1 | RFP 0 | Mac-Tum Doublet |
| DAPI 0 | GFP 1 | DAPI 2 | RFP 0 | Removed |
| DAPI 0 | GFP 1 | DAPI 2 | RFP 1 | Removed |
| DAPI 0 | GFP 1 | DAPI 2 | RFP 2 | Removed |
| DAPI 1 | GFP 1 | DAPI 1 | RFP 1 | Removed |
| DAPI 1 | GFP 1 | DAPI 0 | RFP 1 | Removed |
| DAPI 1 | GFP 1 | DAPI 1 | RFP 0 | Macrophage |
| DAPI 1 | GFP 1 | DAPI 2 | RFP 0 | Mac-Tum Doublet |
| DAPI 1 | GFP 1 | DAPI 2 | RFP 1 | Mac-Tum Doublet |
| DAPI 1 | GFP 1 | DAPI 2 | RFP 2 | Mac-Tum Doublet |

#### Library Quantification

All libraries were benchmarked using the Qubit dsDNA High Sensitivity assay (ThermoFisher Q32851). Concentrated libraries were diluted down to approximately 2 ng/µL. Subsequently, the molarity of the DNA was confirmed via the Agilent Tapestation 4150 D500 tape (Agilent 5067-5592).

#### Library Sequencing Parameters

All library preparations were sequenced using standard chemistry and following standard protocols on the Illumina NextSeq2000 for 650pm standard flow cells and 488pm for X-LEAP flow cells.

The *in vitro* 10x scRNA library was sequenced as paired-end with 28 cycles for read 1, 90 cycles for read 2, 10 cycles for index 1, and 10 cycles for index 2 utilizing a P2-200 X-LEAP flow cell (Illumina Inc. 20100986).

The Artificial Doublet library was sequenced as paired-end with 75 cycles for read 1, 75 cycles for read 2, 8 cycles for index 1, and 8 cycles for index 2 utilizing P2-200 (Illumina Inc. 20046812). The 10x scRNA patient sample libraries were sequenced as paired-end with 28 cycles for read 1, 90 cycles for read 2, 10 cycles for index 1, and 10 cycles for index 2 utilizing P2-100 ((Illumina Inc. 20046811).

The SmartSeq scRNA patient libraries were sequenced as paired-end with 100 cycles for read 1, 100 cycles for read 2, 10 cycles for index 1, and 10 cycles for index 2 utilizing P2 -200 and P3-200 X-LEAP flow cells (Illumina Inc. 20100989).

ScaleBio tagmented scATAC libraries were sequenced as paired-end with 85 cycles for read 1, 125 cycles for read 2, 10 cycles for index 1, and 10 cycles for index 2.

### Computational Analysis scRNA & scATAC

#### Primary Sequence Data Processing for 10x Genomics scRNA Libraries

Raw sequence files were processed using CellRanger^107^ count and aligned to either GRCh38 or MM10 to generate matrix, features, and barcodes files for downstream processing in Seurat.

#### Primary Sequence Data Processing for SmartSeq scRNA Libraries

Sequence reads were demultiplexed using unidex (https://github.com/adeylab/unidex) which matches index barcodes to a whitelist. STAR (https://github.com/alexdobin/STAR.git) was utilized to align demultiplexed fastq files to GRCh38. Scitools rmdup (https://github.com/adeylab/scitools) was utilized to remove duplicate reads. The subread (https://github.com/ShiLab-Bioinformatics/subread.git) module was utilized to perform annotations against GRCh38, and the subsequent feature counts were converted to a count matrix using a custom perl script (available through github repository under data availability). This matrix was suitable for downstream analysis in Seurat.

#### Analysis of scRNA Data

Matrices were read into Seurat and used to generate Seurat objects.^90,91^ The data was normalized, variable features were determined and scaled, and dimensions were reduced via PCR using default parameters. Native functions for finding nearest neighbors, clustering and UMAP projection were employed using standard parameters.

After metadata annotation, like experiments were merged into combined Seurat objects, and re-run through prior functions. Elbow plots were utilized to determine reasonable cut-offs for PCA dimensions, typically 10. Expression of various genes of interest were visualized across clusters using feature plot functions and GO term enrichment analysis was conducted using clusterprofiler package.^94^

#### Primary Sequence Data Processing for scATAC Library

Unidex mode iCell8_scale24 was employed for demultiplexing based on the two 10 bp indices, matched to respective index files. Additionally, the first 8bp of read 2 served as the tagmentation index for the ScaleBio indexed Tn5. These 8 bp of Tn5 index were trimmed together with the next 20 bp of mosaic end sequence. The demultiplexed reads were then aligned to the human reference genome hg38 using bwa-mem (v0.7.15-r1140), processed with scitools rmdup, and then filtered down to cell barcodes reaching a minimum unique read count.

#### Analysis of scATAC Data

Aligned, duplicate-filtered bam files were imported into ArchR to generate an arrow file, with sample names added via a separate text file, and compiled into an ArchR project.^108^ Annotations were subsequently imported from a CSV and added as a meta data column in the ArchR project. Iterative LSI, UMAP projections, TSNE projections were performed using default parameters. Track plots were generated for genes of interest and genes in associated pathways. MACS2 was employed for peak calling and identifying marker peaks, which were used for peak enrichment analysis in a pair-wise fashion.

### Statistical Analysis

All statistical analyses were performed using GraphPad Prism (version 10.6.0) or R (version 4.3.2). Statistical tests were selected based on data distribution, sample size, and experimental design as follows: one-way ANOVA with Dunnett’s post hoc test was used for multiple group comparisons with a shared control (tumor vs. each hybrid cell line) in normally distributed datasets (Figure 2D,2F, 4E). Unpaired two-tailed Student’s t-tests were used for comparisons between two groups in normally distributed data (Figure 2G; 5I). Tukey’s multiple comparisons was used for *in vivo* tumor growth analysis (Figure 2I) and a mixed-effects model with uncorrected Fisher’s LSD was used to compare *in vivo* tumor growth size increase during RUNX1 inhibitor treatment (Figure 5K). For non-normally distributed data, a Kruskal-Wallis test followed by pairwise Wilcoxon rank-sum tests was used for group comparisons (tumor vs. each hybrid cell line), with Benjamini-Hochberg false discovery rate (FDR) correction applied for multiple testing (Figure 4F; 5E; 6G). A Mann-Whitney test was used to compare the number of H11 cells in circulation after RUNX1 treatment (Figure 5L). A two-way ANOVA with a generalized linear mixed model (GLMM) with a beta distribution was employed for proportional data (Figure 6A). Pairwise Wilcoxon rank-sum tests were performed for selected nonparametric comparisons, with Cliff’s delta used to quantify effect sizes (Figure 7I). Results are reported as mean ± standard deviation (SD). All violin plots, UMAPs, and dot plots were generated using the ggplot2 and Seurat packages in R. Statistical significance was defined as *p* < 0.05.

