## Supplementary Figures for "Cell fusion reprograms tumor cells and promotes RUNX1-mediated invasion and dissemination in colorectal cancer"

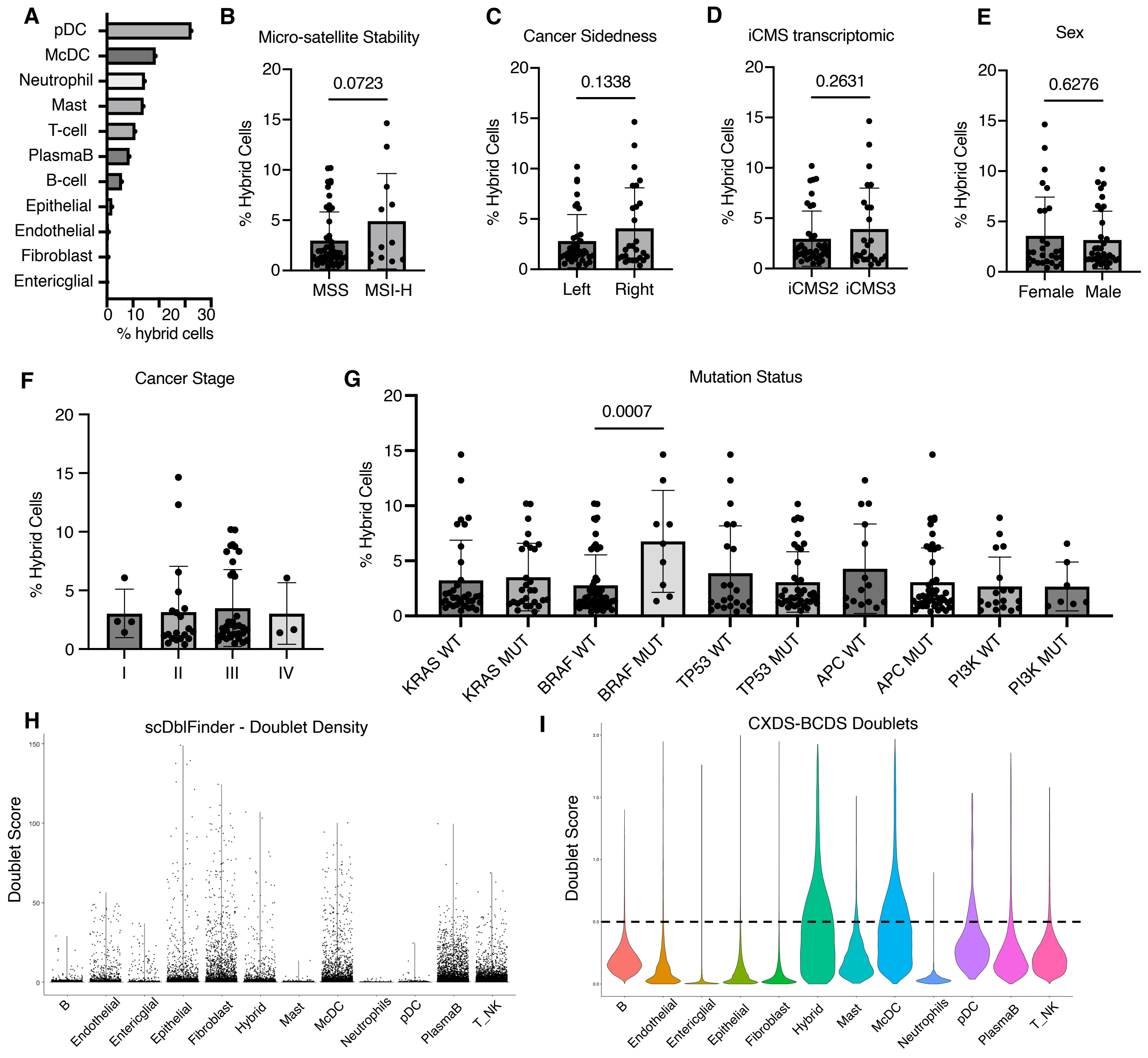
A
pDC
McDC
Neutrophil
Mast
T-cell
PlasmaB
B-cell
Epithelial
Endothelial
Fibroblast
Entericglial
% hybrid cells
B
Micro-satellite Stability
0.0723
% Hybrid Cells
MSS
MSI-H
C
Cancer Sidedness
0.1338
Left
Right
D
iCMS transcriptomic
0.2631
iCMS2
iCMS3
E
Sex
0.6276
Female
Male
F
Cancer Stage
I
II
III
IV
G
Mutation Status
0.0007
KRAS WT
KRAS MUT
BRAF WT
BRAF MUT
TP53 WT
TP53 MUT
APC WT
APC MUT
PI3K WT
PI3K MUT
H
scDblFinder - Doublet Density
Doublet Score
B
Endothelial
Entericglial
Epithelial
Fibroblast
Hybrid
Mast
McDC
Neutrophils
pDC
PlasmaB
T_NK
I
CXDS-BCDS Doublets
Doublet Score
B
Endothelial
Entericglial
Epithelial
Fibroblast
Hybrid
Mast
McDC
Neutrophils
pDC
PlasmaB
T_NK

**Supplemental Figure 1. Clinical and molecular features associated with hybrid cell abundance in CRC.** A) Bar graph showing the proportion of hybrid cells across annotated immune and stromal cell types, including pDCs, mDCs, neutrophils, mast cells, T cells, plasma cells, B cells, epithelial, endothelial, fibroblast, and enteroglial populations. B–E) Comparison of hybrid cell frequency across patient subgroups stratified by clinical and molecular characteristics, including (B) microsatellite stability (MSS vs MSI-H), C) tumor sidedness (left vs right colon), D) iCMS transcriptomic subtype (iCMS2 vs iCMS3), and E) sex. F) Hybrid cell abundance across cancer stage (I–IV). G) Association between hybrid cell frequency and mutation status of commonly altered genes in CRC including KRAS, BRAF, TP53, APC, and PI3K. H) Distribution of doublet scores calculated by scDblFinder across all samples. I) Violin plots showing doublet scores from CXDS-BCDS combined method across major cell types. Dashed line indicates the doublet threshold. Statistical significance determined using unpaired two-tailed t-tests.

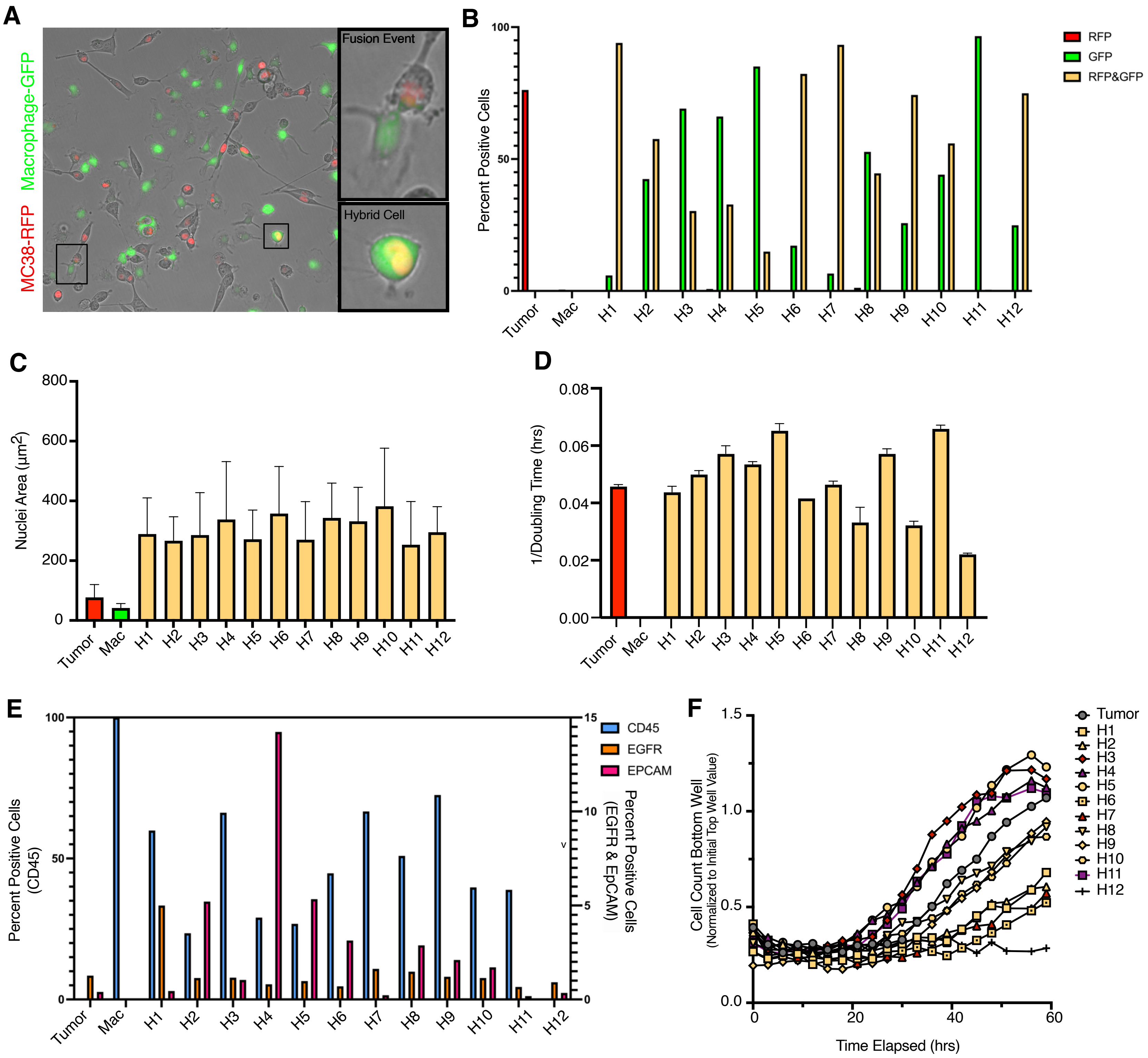
A
MC38-RFP Macrophage-GFP
Fusion Event
Hybrid Cell
B
Percent Positive Cells
RFP
GFP
RFP&GFP
Tumor
Mac
H1
H2
H3
H4
H5
H6
H7
H8
H9
H10
H11
H12
C
Nuclei Area (μm2)
D
1/Doubling Time (hrs)
E
Percent Positive Cells (CD45)
Percent Positive Cells (EGFR & EpCAM)
CD45
EGFR
EPCAM
F
Cell Count Bottom Well
(Normalized to Initial Top Well Value)
Time Elapsed (hrs)
Tumor

**Supplemental Figure 2. Phenotypic heterogeneity of hybrid single cell clones (H1-H12).** A) Images showing MC38-RFP (Tumor) and primary bone marrow derived macrophages-GFP (Mac) on day 3 of co-culture with fusion event and hybrid cell magnified insets. B) Bar graph showing percent positive cells for RFP, GFP, or dual RFP+GFP+ expression across hybrid clones (H1–H12) compared to unfused tumor cells. C) Quantification of nuclear size (area, $\mu m^2$) for each hybrid clone and parental MC38 tumor and macrophage cells. D) Proliferation rates measured by phase confluence imaging over 3 days. E) Surface marker antibody expression across hybrid clones measured by flow cytometry. F) Trans-well chemotaxis assay of hybrid single cell clones and MC38 tumor cells, measured by phase object count over a 60-hour time course where each symbol represents a different clone.

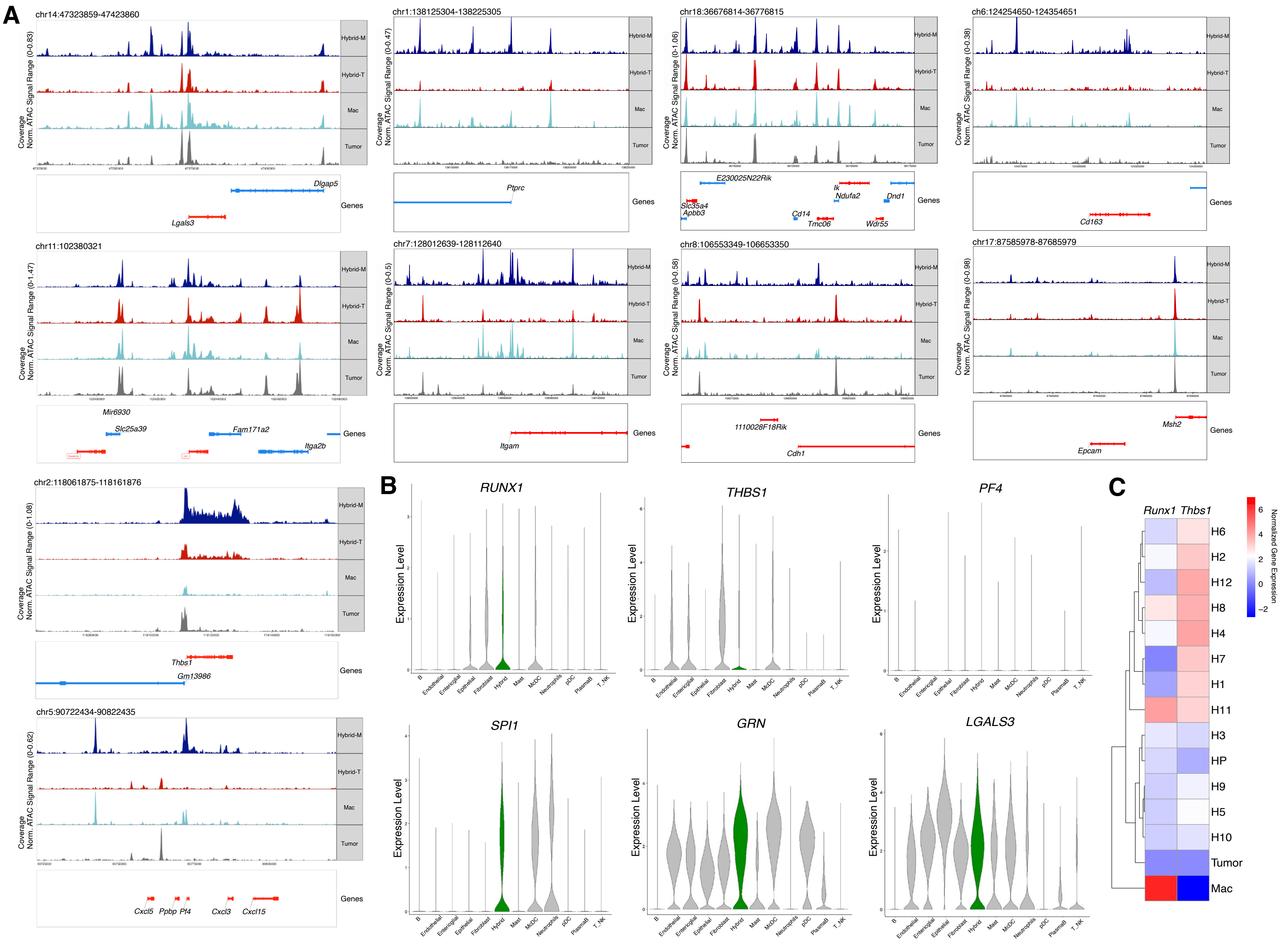
A
chr14:47323859-47423860
chr1:138125304-138225305
chr18:36676814-36776815
chr6:124254650-124354651
Coverage
Norm. ATAC Signal Range (0-0.83)
Norm. ATAC Signal Range (0-0.47)
Norm. ATAC Signal Range (0-1.06)
Norm. ATAC Signal Range (0-0.38)
Hybrid-M
Hybrid-T
Mac
Tumor
Genes
Dlgap5
Lgals3
Ptprc
E230025N22Rik
Ik
Ndufa2
Dnd1
Slc35a4
Apbb3
Cd14
Tmc06
Wdr55
Cd163
chr11:102380321
chr7:128012639-128112640
chr8:106553349-106653350
chr17:87585978-87685979
Norm. ATAC Signal Range (0-1.47)
Norm. ATAC Signal Range (0-0.5)
Norm. ATAC Signal Range (0-0.58)
Norm. ATAC Signal Range (0-0.98)
Mir6930
Slc25a39
Fam171a2
Itga2b
Itgam
1110028F18Rik
Cdh1
Msh2
Epcam
chr2:118061875-118161876
Norm. ATAC Signal Range (0-1.08)
Thbs1
Gm13986
chr5:90722434-90822435
Norm. ATAC Signal Range (0-0.62)
Cxcl5
Ppbp
Pf4
Cxcl3
Cxcl15
B
RUNX1
THBS1
PF4
SPI1
GRN
LGALS3
Expression Level
B
Endothelial
Enteroglial
Epithelial
Fibroblast
Hybrid
Mast
MoDC
Neutrophils
pDC
PlasmaB
T_NK
C
Runx1
Thbs1
H6
H2
H12
H8
H4
H7
H1
H11
H3
HP
H9
H5
H10
Tumor
Mac
Normalized Gene Expression
6
4
2
0
−2

**Supplemental Figure 3. Chromatin accessibility and expression of RUNX1 pathway genes in hybrid cells.** A) ATACseq track plots of hybrid-M, hybrid-T, tumor and macrophage cell populations for additional macrophage and epithelial genes including Ptprc (Cd45), Cd14, Cd163, Itgam, Cdh1 (Ecad), Epcam, and Runx1 pathway genes: Lgals3, Thbs1, Grn and Pf4. B) Runx1 pathway gene expression in CRC human hybrids from scRNA-seq studies introduced in Figure 1, and C) *Runx1* and *Thbs1* expression in MC38-macrophage fusion hybrid single cell clones measured by qRT-PCR.

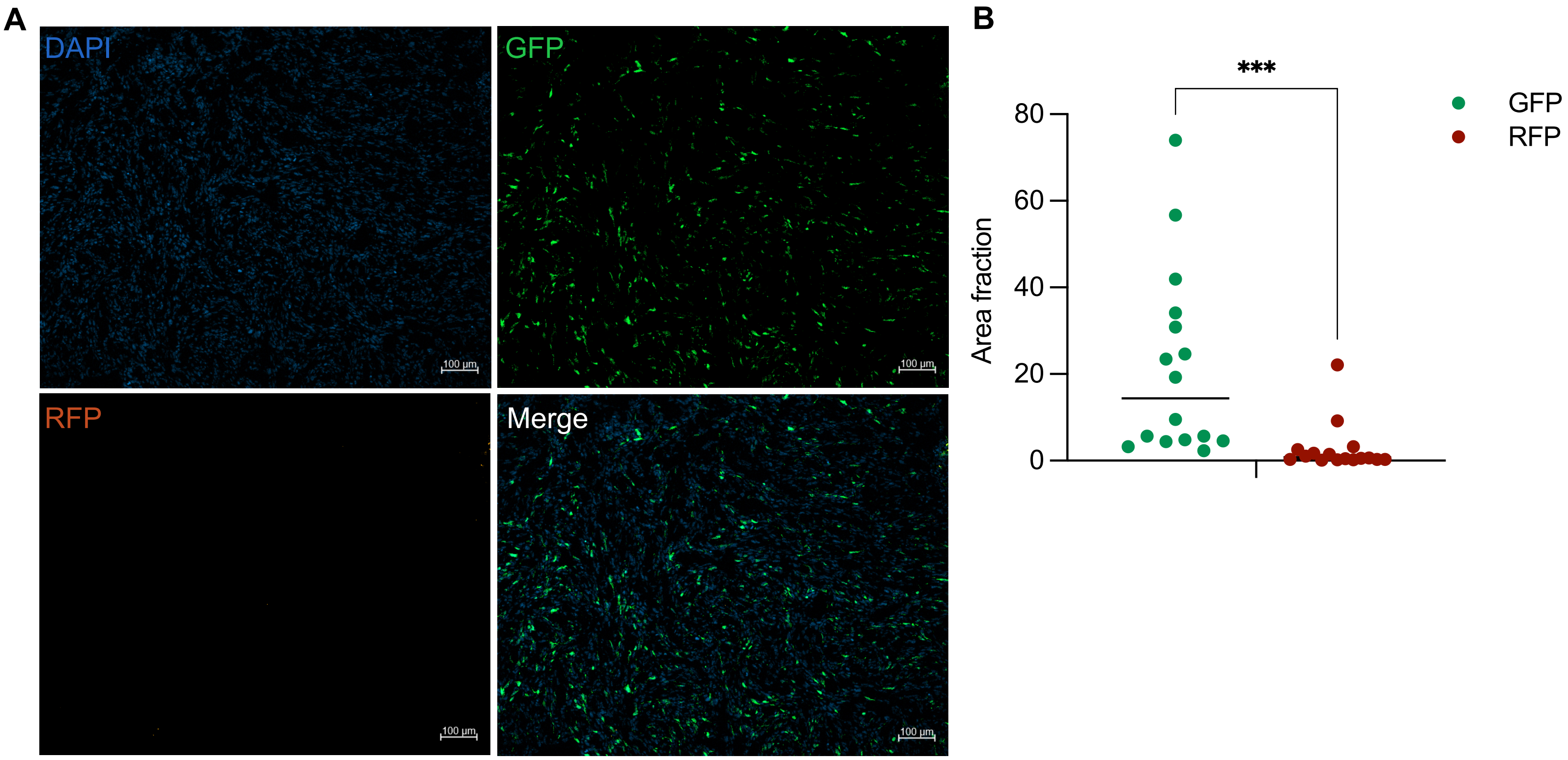
A
DAPI
GFP
RFP
Merge
100 µm
100 µm
100 µm
100 µm
B
***
80
60
40
20
0
Area fraction
GFP
RFP

**Supplemental Figure 4. Fluorescence reporter signal quantification in H11 hybrid cell tumors.** A) Representative immunofluorescence images of GFP and RFP reporter signals in H11 mouse tumor. B) Quantification of GFP and RFP reporter signal as area fraction above threshold in outer (n=2) and inner (n=2) regions from H11 tumors (n=4). Statistical significance determined by paired t-test (***$p$ <0.001).

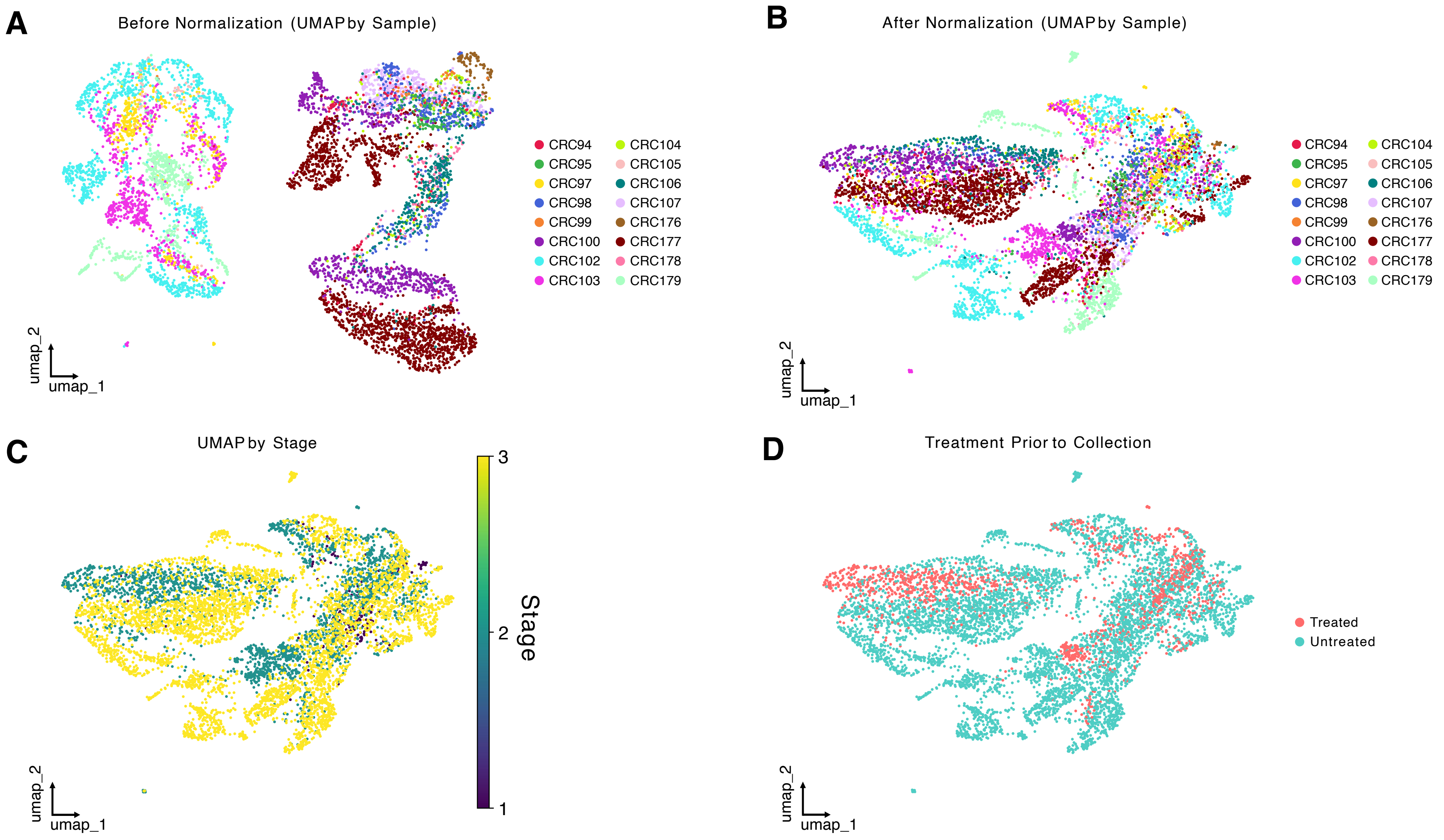
A
Before Normalization (UMAP by Sample)
B
After Normalization (UMAP by Sample)
CRC94
CRC95
CRC97
CRC98
CRC99
CRC100
CRC102
CRC103
CRC104
CRC105
CRC106
CRC107
CRC176
CRC177
CRC178
CRC179
umap_1
umap_2
C
UMAP by Stage
Stage
3
2
1
D
Treatment Prior to Collection
Treated
Untreated

**Supplemental Figure 5. Cyclic immunofluorescence analysis of CHCs in CRC.** A) UMAP plot of cyclic immunofluorescence (cyCIF) data before normalization colored by patient ID. B) UMAP plot of cyCIF data after normalization colored by patient ID. C) UMAP plot of cyCIF data colored by stage. D) UMAP plot of cyCIF data colored by treatment status prior to sample collection.

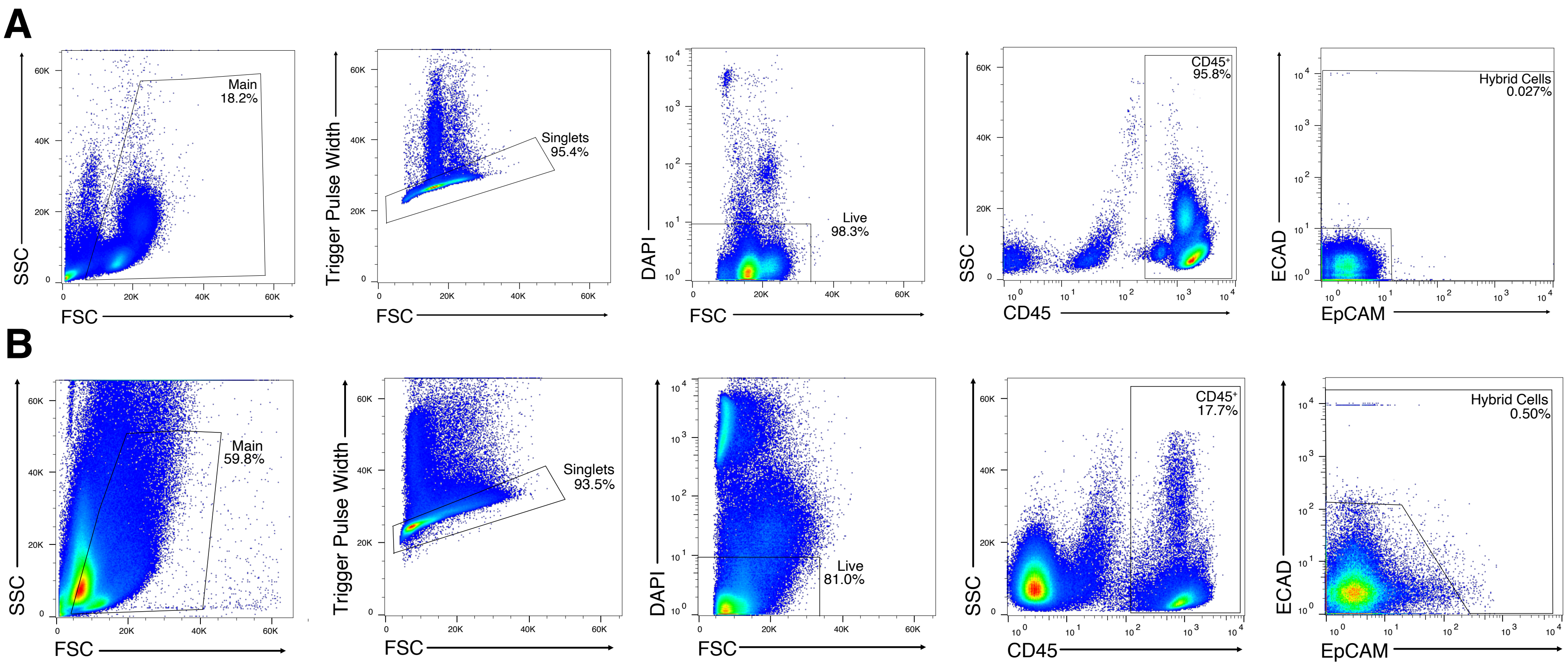
A
B
SSC
FSC
Main
18.2%
Trigger Pulse Width
Singlets
95.4%
DAPI
Live
98.3%
CD45⁺
95.8%
CD45
ECAD
EpCAM
Hybrid Cells
0.027%
Main
59.8%
Singlets
93.5%
Live
81.0%
CD45⁺
17.7%
Hybrid Cells
0.50%

**Supplemental Figure 6. Flow cytometry gating strategy for hybrid cells isolated from CRC primary tumor and peripheral blood.** Sequential gating strategy for hybrid cells isolated from A) peripheral blood (CHCs) and B) primary CRC tumors (TrHCs). Gating from left to right includes debris, doublet exclusion, live cell gating, identification of $CD45^+$ cells, and identification of $ECAD^+$ and/or $EpCAM^+$.

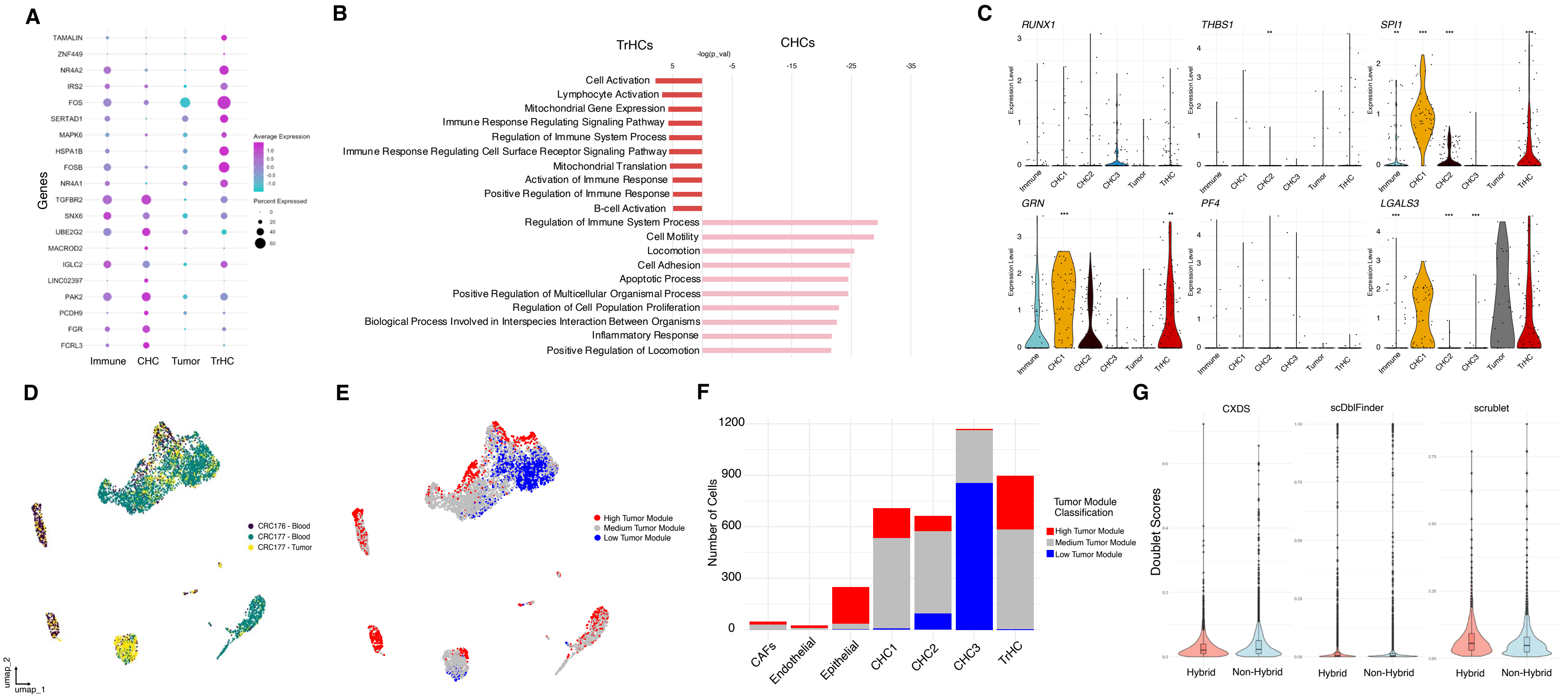
A
TAMALIN
ZNF449
NR4A2
IRS2
FOS
SERTAD1
MAPK6
HSPA1B
FOSB
NR4A1
TGFBR2
SNX6
UBE2G2
MACROD2
IGLC2
LINC02397
PAK2
PCDH9
FGR
FCRL3
Genes
Immune
CHC
Tumor
TrHC
Average Expression
Percent Expressed
B
TrHCs
CHCs
-log(p_val)
Cell Activation
Lymphocyte Activation
Mitochondrial Gene Expression
Immune Response Regulating Signaling Pathway
Regulation of Immune System Process
Immune Response Regulating Cell Surface Receptor Signaling Pathway
Mitochondrial Translation
Activation of Immune Response
Positive Regulation of Immune Response
B-cell Activation
Regulation of Immune System Process
Cell Motility
Locomotion
Cell Adhesion
Apoptotic Process
Positive Regulation of Multicellular Organismal Process
Regulation of Cell Population Proliferation
Biological Process Involved in Interspecies Interaction Between Organisms
Inflammatory Response
Positive Regulation of Locomotion
C
RUNX1
THBS1
SPI1
GRN
PF4
LGALS3
Expression Level
Immune
CHC1
CHC2
CHC3
Tumor
TrHC
D
CRC176 - Blood
CRC177 - Blood
CRC177 - Tumor
umap_2
umap_1
E
High Tumor Module
Medium Tumor Module
Low Tumor Module
F
Number of Cells
1200
900
600
300
0
CAFs
Endothelial
Epithelial
CHC1
CHC2
CHC3
TrHC
Tumor Module Classification
High Tumor Module
Medium Tumor Module
Low Tumor Module
G
Doublet Scores
CXDS
scDblFinder
scrublet
Hybrid
Non-Hybrid

**Supplemental Figure 7. Additional information from patient matched CHC and TrHC scRNA-sequencing studies.** A) Dot plot of top differentially expressed genes between circulating hybrid cells (CHCs) and tumor resident hybrid cells (TrHCs) from patient matched CRC primary tumor and peripheral blood scRNA SMART-seq dataset. B) Gene set enrichment analysis of GO biological processes comparing CHCs to TrHCs. C) Violin plots of Runx1 pathway genes including *RUNX1, THBS1, SPI1, GRN, PF4,* and *LGALS3*, across immune, tumor, CHC, and TrHC populations from scRNA-seq SMART-seq studies. D) Harmony-integrated UMAP of 10X scRNA-seq data from patient-matched CRC tumor and peripheral blood specimens. E) UMAP of 10X scRNA-seq dataset colored by tumor module score classification (derived from tumor cell controls from SMART-seq dataset). F) Quantification of tumor module score classification by cell type. G) Doublet detection analysis using CXDS, scDblFinder, and scrublet across hybrid and non-hybrid populations from 10X scRNA-seq dataset.
